# The *Mycobacterium smegmatis bd*-II terminal oxidase employs a carboxylate shift mechanism

**DOI:** 10.1101/2025.06.23.660515

**Authors:** Terezia Kovalova, Mateusz Janczak, Ana P. Gamiz-Hernandez, Daniel Lundin, Soni Sharma, Johanna Vilhjálmsdóttir, Dan Sjöstrand, Ville R. I. Kaila, Martin Högbom, Pia Ädelroth

**Author notes:** corresponding authors: Ville R. I. Kaila, Martin Högbom, Pia Ädelroth. equal contribution (Terezia Kovalova, Mateusz Janczak, Ana P. Gamiz-Hernandez).

## Abstract

Cytochrome *bd* is a terminal oxidase expressed under low oxygen conditions and central for the survival of many pathogens. Here we characterise the first qOR-2 type *bd* oxidase, the cyt *bd*-II from *Mycobacterium smegmatis*, by combining biochemical studies with cryo-electron microscopy (cryo-EM), and multiscale simulations. By over-expressing the *appCB* operon in its native host, we produce a highly active *bd*-II (*k*_cat_=30 e^-^s^-1^) that together with a high-resolution (2.8 Å) cryo-EM structure and multiscale simulations reveal unique proton pathways and oxygen channels responsible for its function. We propose that O_2_-scavenging activates a pH-dependent molecular switch, involving coordination changes of heme *d* and surrounding bulky residues that regulate substrate access into the active site. Taken together, our findings provide detailed mechanistic insight of qOR-2 type *bd* oxidases, and a basis for understanding the evolution of the superfamily.

---

The respiratory chain of mitochondria and prokaryotes catalyse electron transfer via membrane-bound protein complexes that generate an electrochemical proton gradient (or proton motive force, PMF), driving ATP synthesis. The final step of bacterial aerobic respiration, the reduction of oxygen to water, is catalysed by both heme-copper oxidases (HCuOs) and cytochrome *bd* oxidases^1,2^. HCuOs comprise a large diverse superfamily, present across all domains of life, comprising three major subfamilies A-C, with the canonical A-type exemplified by the O_2_-reducing, proton-pumping mitochondrial cytochrome *c* oxidase (Complex IV). In contrast, the cyt *bd* quinol oxidases are evolutionarily unrelated to the HCuOs and only found in prokaryotes, mostly bacteria^3^.

The cyt *bd*s have a higher O_2_ affinity than HCuOs^4^ and are less sensitive to common HCuO inhibitors such as cyanide and NO^5^. These properties endow the cyt *bd*-harboring bacteria with important physiological functions, such as protection of O_2_-sensitive enzymes (such as nitrogenase) against degradation and the capability to colonize microaerophilic environments^6^. The cyt *bd* family are often crucial for pathogenicity of bacteria, making them promising drug targets^7^. This property is presumably related to the involvement of the cyt *bd* in protection against the oxidative and nitrosative stress defenses of the host^5,8^, see also Refs.^5,9,10^.

All characterised cyt *bd*s share two core subunits, CydA (AppC) and CydB (AppB), that have similar architecture, comprising two transmembrane four-helix bundles and one peripheral helix each (Fig. 1a), with the cofactors heme *b*^558^, heme *b*^595^, and heme *d* located within CydA (AppC). O_2_ binds and is reduced at heme *d*, whereas the membrane-soluble electron donor quinol (Q) binds close to six-coordinated heme *b*^558^ in the so-called Q-loop, near the periplasmic surface. The Q-loop shows variation in length, with both short and long Q-loops present in various *bd*s. Heme *b*^595^ is a penta-coordinated high-spin heme, but generally not presumed to participate in O_2_ binding^9^. Protons are transferred from the cytoplasmic side to the heme *d* active site for reduction of O_2_, whereas the quinol oxidation releases protons to the periplasm, rendering the reaction electrogenic. However, these reactions presumably do not couple to proton pumping, suggesting that the cyt *bd*s conserve less of the free energy than the HCuOs^6^.

**Fig. 1.**
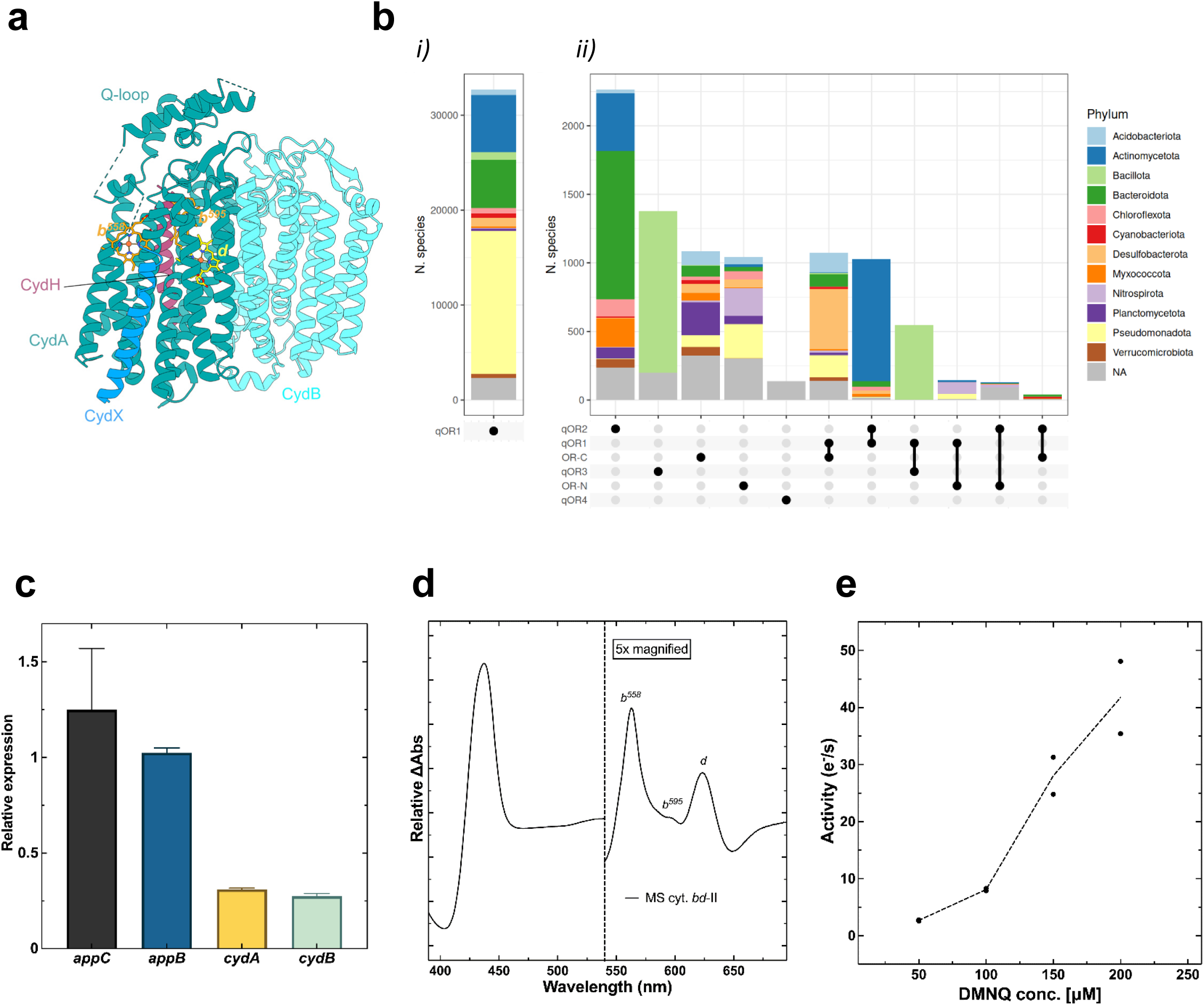
Gene expression, protein purification and characterization of the cyt *bd*-II from *M. smegmatis*. **a,** General overview of a cyt *bd* oxidase (here *E. coli* cyt *bd*-I, PDB ID: 6RKO^19^). Two core subunits are depicted in dark blue (CydA, containing Q-loop) and teal (CydB) with the three heme cofactors depicted in orange (heme *b^558^* and *b^595^*) and yellow (heme *d*). Accessory subunits depicted in blue (CydX) and purple (CydH). **b**, Species encoding combinations of different cyt *bd* subfamilies, *i*) the twelve most numerous phyla encoding *only* qOR1, *ii)* the eleven most common combinations of any of the other *bd* subfamilies. **c**, Real-time RT-PCR mRNA expression for *appCB* compared to *cydAB* in *M. smegmatis* during fast and slow-growing conditions. **d**, Dithionite-reduced *minus* air-oxidized spectra of the *Ms* cyt *bd*-II. **e**, Oxygen reducing/menaquinol oxidizing activity of *Ms* cyt *bd-*II with varying concentrations of reduced DMNQ (50-200 μM) added. Data points shown for each individual measurement, and the line drawn through the average.

In Actinomycetota (formerly Actinobacteria) such as *Mycobacterium smegmatis* and *M. tuberculosis*, the respiratory chain is branched, with the menaquinol oxidation catalysed by either the *bcc*-*aa*_3_ (III_2_/IV_2_) supercomplex or by the cyt *bd* branch^11^. *M. tuberculosis* harbours one cyt *bd*, encoded by the *cydAB* operon^11,12^, shown to be important for the survival of the pathogen. Dual targeting of the *bcc-aa_3_* branch and the *bd* branch has been recognized as an effective strategy for drug development against tuberculosis^13,14^.

Many Actinomycetota contain several genes encoding for cyt *bds*^3^ and *M. smegmatis* harbours both *cydAB genes* (encoding the *Ms bd*-I), homologous to the *M. tuberculosis cydAB*, shown to be important for microaerobic respiration in *M. smegmatis*^11^, similarly to its role in other organisms. The *Ms bd*-I has been biochemically and structurally characterised^15,16^, whilst in contrast, the *appCB* genes, found next to the *cydAB* operon have a low sequence identity to CydAB (∼25% for CydA) and remain uncharacterised. The *appCB* genes are transcribed ^17^ and upregulated under conditions where the *bcc-aa_3_* branch is inactivated and no functional CydAB is present^17,18^, but the functional role for *Ms* AppCB *(bd*-II) remains poorly understood.

The cyt *bd* superfamily was recently classified using an extensive set of CydA sequences, yielding a division of the O_2_-reducing cyt *bd*s into four subfamilies, qOR-1 to qOR-4a^3^. Out of these, qOR-1s, to which the *E. coli* cyt *bd*-I (and *bd*-II) belongs, is well-characterised. The *Ms bd*-I (cydAB) is also a member of the qOR-1 class, whereas the *Ms bd*-II (AppCB) is a part of the qOR-2 class. The qOR-2 subfamily is widely distributed in bacterial phyla^3^, and highly prevalent among Actinomycetota. The currently determined structures of the *bd* superfamily are dominated by the qOR-1 class (the *E. coli bd*-I^19,20^ and *bd*-II^21,22^, the *M. tuberculosis*^23^ and *M. smegmatis* cyt *bd*-I, and the *Corynebacterium glutamicum bd*^24^). Moreover, there is one structure from the qOR-3 class (*Geobacillus thermodenitrificans bd*^25^), but currently no member of the qOR-2 group is neither biochemically nor structurally characterised.

Here, we study the first qOR-2, the *Ms bd*-II, by combining biochemical experiments with structure determination, bioinformatic analysis, and multiscale simulations. The isolated *Ms bd-*II complex is active with menaquinol as substrate, and the determined high-resolution structure together with molecular simulations reveal key features of its molecular architecture, including the location of several structural menaquinone molecules, as well as proton and oxygen channels. Our combined findings provide a basis for understanding the mechanistic adaptations of the qOR-2 *bd* family and sheds light on the evolution of the *bd* superfamily.

## Results

### Prevalence and distribution of the qOR-2 *bd* family with focus on Actinomycetota

To explore the distribution of *bd* oxidases, sequences were analysed (see *Methods*) based on data collected in Ref.^3^. The qOR-1 class is by far the most common *bd* subfamily across phyla, with individual species expressing only this type being most common. qOR-2s represent the second most common class of *bd*-oxidases, followed by the qOR-3s, exemplified by the *Geobacillus thermodenitrificans* (*Gt,* of the *Bacillota* phylum) *bd*. A combination of qOR-1 and qOR-2 is commonly found in the Actinomycetota (Fig. 1b, Supplementary Fig. S1), as exemplified by *M. smegmatis*, studied here. Although the *Ms bd*-II of the qOR-2 family is not the only *bd* of this organism, there are several examples of species that harbor only a qOR-2 *bd*, especially other Actinomycetota, such as the *Frankiaceae* family containing the nitrogen-fixing *Frankia spp*. (Supplementary Fig. S1 and Supplementary Table S1), where the *bd* has a possible functional role as an oxygen scavenger. The unique features identified from our sequence alignments for both CydA (AppC) and CydB (AppB) were further used to complement the functional analysis (see below).

### *M. smegmatis* appCB gene expression, protein expression, purification, and characterization

Expression of the *appCB* (*Ms bd*-II) mRNA was compared to *cydAB* (Ms *bd*-I) using real time RT-PCR and showed transcriptional upregulation of the *appCB* genes at lower O_2_ conditions (Fig. 1c). The *appCB* genes were recombinantly expressed in *M. smegmatis*, membranes were solubilised with the detergent lauryl maltose neopentyl glycol (LMNG), and the *Ms bd*-II was purified to homogeneity (Supplementary Fig. S2). SDS-PAGE and mass-spectrometric analysis support the expression of the AppC (∼28% coverage) and AppB (∼12%) proteins, and an additional peptide belonging to an unknown protein (Supplementary Fig. S2). Spectral analysis of the *Ms bd*-II (AppCB) protein shows heme peaks consistent with a heme *b*^558^, heme *b*^595^, and heme *d* (Fig. 1d), similar to previously characterized *cyt bd*s of the qOR-1 and qOR-3 classes ^16,22,23,25^. However, the characteristic heme *d*^2+^-O_2_ peak at ∼650 nm in the ‘as isolated’ spectrum is smaller in the *Ms bd*-II than in the well-characterized *E. coli bd*-I (Supplementary Fig. S3). The *Ms bd*-II is active in oxygen reduction assays using 1,4-naphthoquinone (DMNQ) as a substrate, with a *k*_cat_ of about 30 e^-^ s^-1^ at 150 µM DMNQ (Fig. 1e), which is in the same range as that of *Ms bd*-I (cydAB)^16^.

### Molecular architecture of the *M. smegmatis* cyt *bd*-II oxidase

The structure of the *Ms bd*-II was determined using cryo-EM to an overall resolution of 2.8 Å (Supplementary Fig. S4 and Supplementary Table S2). The protein complex consists of two subunits, AppC and AppB (Fig. 2). The overall fold is nearly identical in the two subunits, with nine transmembrane helixes, organised into a bundle (Fig. 2). The structure has an overall resemblance to previously solved *bd* structures^10^ (Supplementary Figs. S5 and S10), but revealing key functional difference unique for the qOR-2 class (see below). There is no density for the additional peptide observed in mass spectrometry, or for homologues of either of the small subunits CydH (found in *Ec bd*-I^19^) or CydX (CydS/AppX, found in several *bd* oxidases^19,25^). CydH in *Ec bd*-I (Fig. 1a) binds at the interface of helix 1 and 8 of CydA (AppC), where it blocks access to the heme *b*^595^. In the *Ms bd*-II map, we observe density that could be interpreted as lipids or detergents at this location (Supplementary Fig. S6). The other qOR-1 structures from mycobacteria also lack additional small subunits^16,23,24^.

**Fig. 2.**
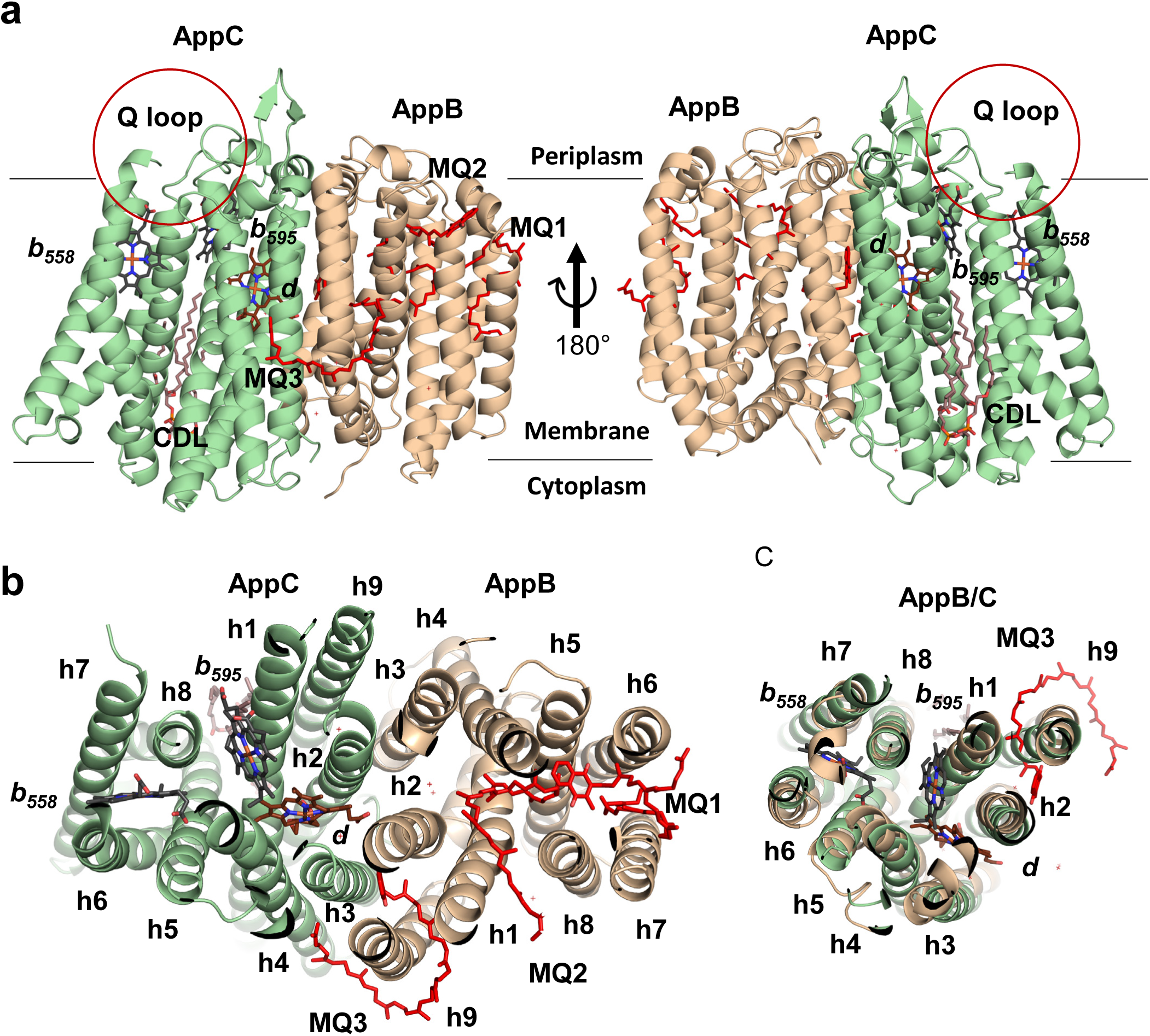
Structure of cyt *bd*-II oxidase from *M. smegmatis*. **a**, The protein complex with individual subunits shown in different colours (AppC in green, AppB in orange), and their relative position within the membrane. Binding sites of hemes (*b^558^*, *b^595^*, and *d*), three menaquinones, and a cardiolipin molecule are marked. Position of the disorder Q loop in the periplasmic side is marked with a red circle. **b**, Top view of the bd-II complex with individual helices numbered and position of the ligands marked. **c**, Superposition of AppC and AppB show fold conservation, and point towards evolution from the same gene.

### Structure of redox-active heme cofactors

The *Ms bd*-II shows a triangular conformation of the hemes, with the same positioning of the individual hemes as observed in *E. coli bd*-I ^19^ and other qOR-1 type *bd*s (Fig. 3, Supplementary Fig. S6). We thus find no evidence for the ‘switch’ between heme *d* and heme *b*^595^, as observed for the *G. thermodenitrificans bd*^25^ of the qOR-3 class. The heme *b*^558^ is located between h5-h6 and h7-h8, and ligated by H197^C^ of h5 and M336^C^ of h7. The highly conserved W385^C^ (Supplementary Fig. S9) is positioned between heme *b*^558^ and heme *b*^595^ and could increase electronic couplings between the cofactors. The penta-coordinated heme *b*^595^ is closest to the periplasmic surface amongst the redox cofactors, between h1 and h8, and ligated by E389^C^ of h8. These helices open up a cavity that could serve as an O_2_ channel (Fig. 4, see below). In the qOR-1 class, the heme *b*^595^ environment also contains a phenylalanine residue (F11^A^ in *Ec bd*-I) that was suggested^19^ to be involved in blocking ligand access (Supplementary Fig. S7). However, this residue is replaced by a methionine (M26^C^) in the *Ms bd*-II and in all other qOR-2 type *bd*s (Supplementary Fig. S9), suggesting that heme *b*^595^ could have a more versatile role.

**Fig. 3.**
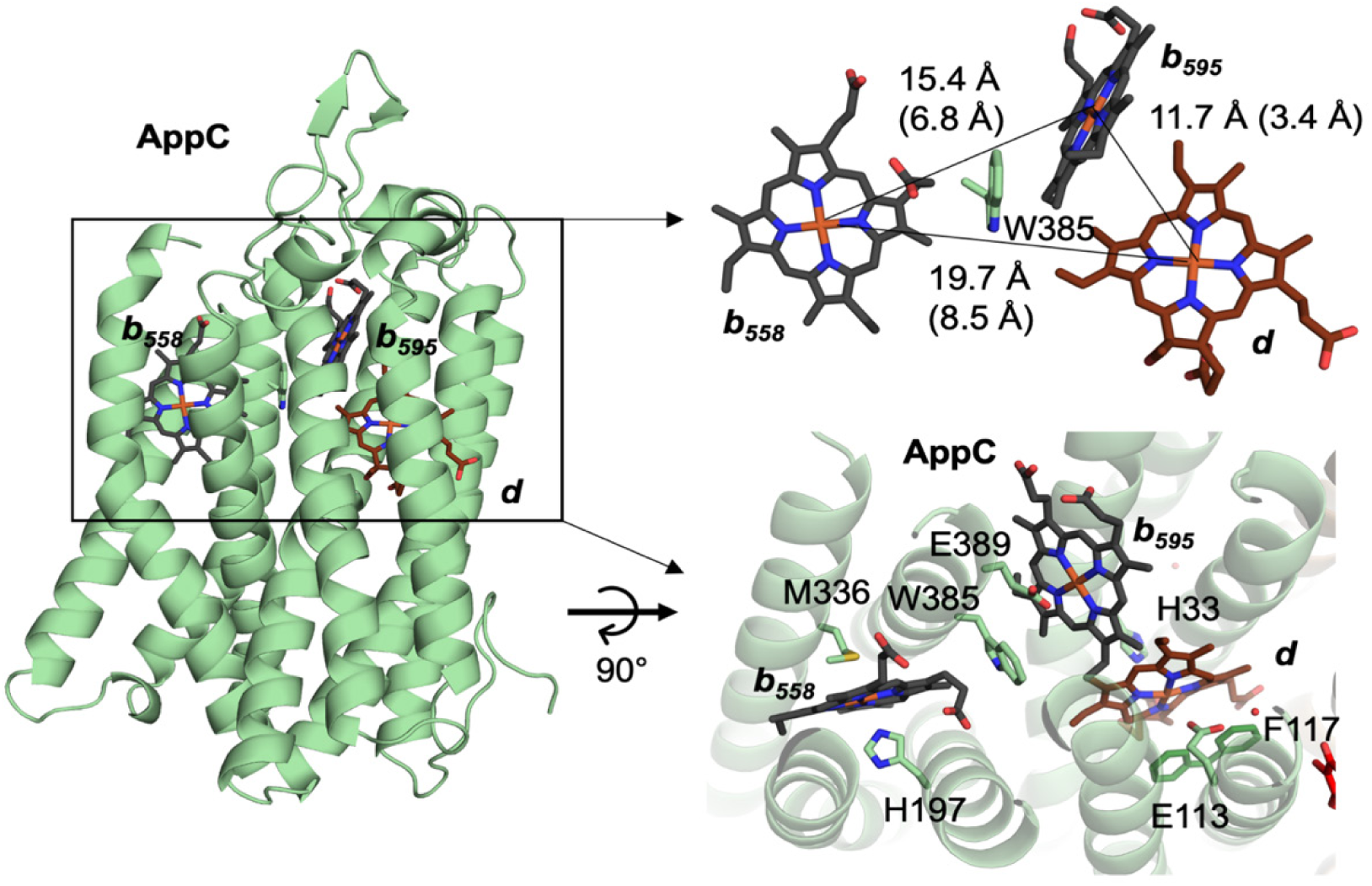
Organisation of heme cofactors in *bd*-II. Position of the individual hemes marked within AppC. Triangular composition of the hemes with distances between individual Fe centres and *edge-to-edge* distances in the brackets. Ligands of individual hemes are marked within AppC. The two conformations of the F117^C^ are indicated.

**Fig. 4.**
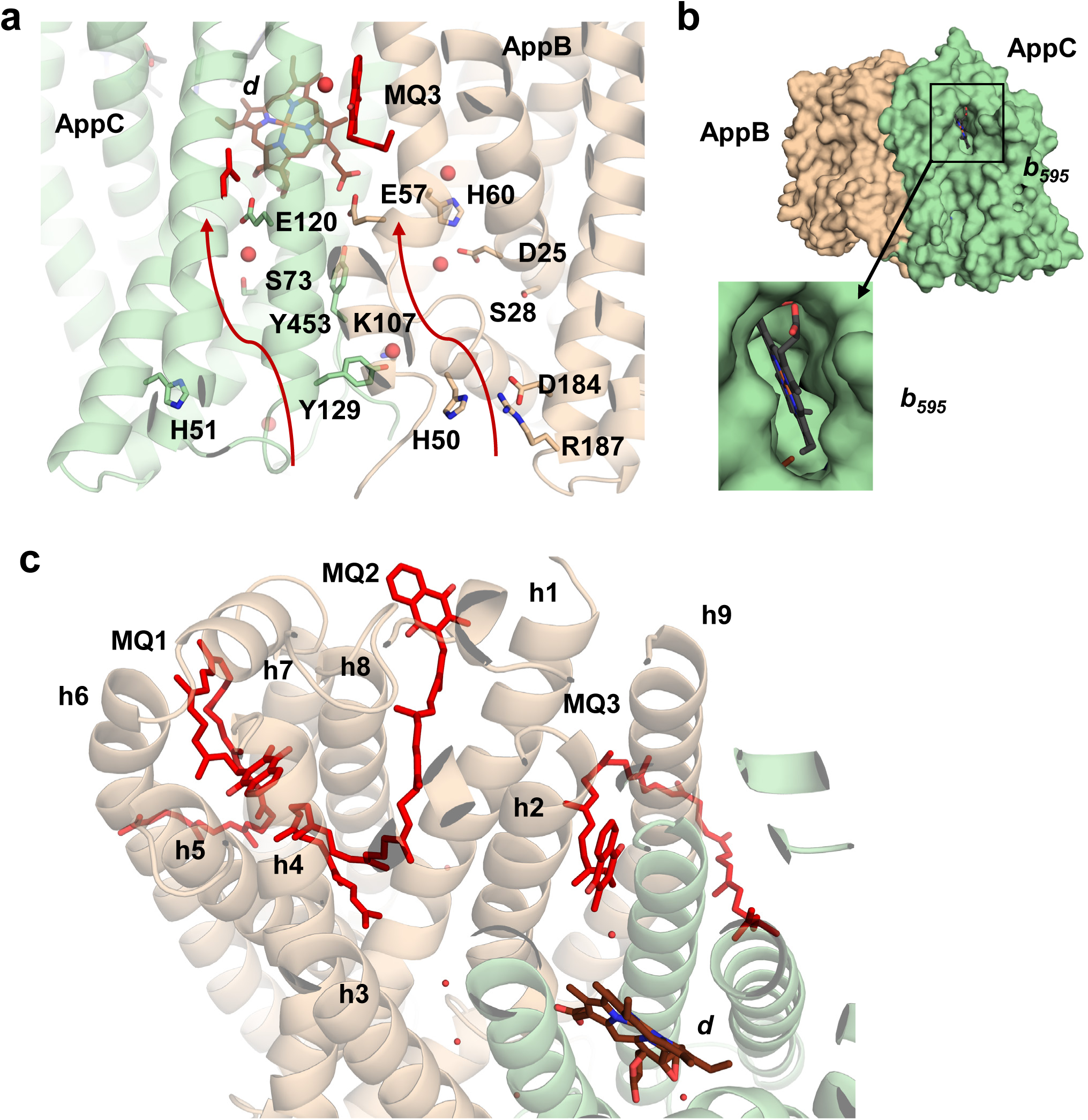
Putative proton pathways and structural MQ binding sites. **a**, Water molecules observed in cryo-EM map are represented as red spheres. Residues forming possible proton channels (red arrows), leading from cytoplasmic side to heme *d* are numbered and represented as sticks. **b**, Surface representation of the protein complex with the open cavity around heme *b*^595^ is highlighted. **c**, Binding sites of the additional menaquinones observed in AppB.

Heme *d* forms the likely O_2_-reduction site of *Ms bd*-II. It is ligated by H33^C^ of h1 and E113^C^ of h2 (Fig. 3 and Supplementary Fig. S8), and positioned at the centre of the membrane within the heme triangle. All heme groups are located in close proximity of each other, with e*dge-to-edge* distances of 3.4 Å (11.7 Å *centre-to-centre*) for heme *b*^595^ – heme *d*; 6.8 Å (15.4 Å *centre-to-centre*) for heme *b*^558^–heme *b*^595^; and 8.5 Å (19.7 Å *centre-to-centre*) for heme *b*^558^–heme *d*, thus providing a structurally efficient electronic wiring between the three redox cofactors. Interestingly, F117^C^, located around 3.4 Å from heme *d*, has two distinct conformations with partial occupancy (Fig. 3). The switch between these states could potentially regulate the access of substrate oxygen and/or protons to the active site, as revealed by our molecular dynamics (MD) simulations (see below).

### Structural MQ binding sites and ligands

We observe three menaquinones bound in the cryo-EM map, with possible structural roles. Two of these sites are positioned within the AppB subunit, whereas the third site is located at the interface of AppB and AppC, near heme *d* (Figs. 2b, 4c). MQ_1_ is located between h5-6 and h7-8, with the aromatic headgroup buried between the helixes, forming contacts with W275^B^, V164^B^, L168^B^, with the hydrophobic tail of MQ_2_ located 3.5 Å from L168^B^. In addition to the contact-forming residues, the MQ_1_ cavity consists of mostly hydrophobic residues (L238^B^, L230^B^, L226^B^, V219^B^, V271^B^, A137^B^, G138^B^, G165^B^). For a comparison to quinone binding sites in other *bd* structures, see Supplementary Fig. S11. Similar non-catalytic, “structural” MQ sites were also previously resolved in the *M. smegmatis* respiratory CIII_2_CIV_2_ supercomplex^26–28^. The hydrophobic tail of MQ_1_ is twisted in a double-loop pointing into the detergent micelle (Fig. 2c).

MQ_2_ is located next to MQ_1_, but inserted tail-first within the complex, with the aromatic headgroup bound to the protein surface, and facing the detergent micelle. The isoprenoid tail of MQ_2_ is twisted in a similar double-loop-like fashion as MQ_1_, coiling around h1, h5, and h8 of AppB. If the AppC and AppB subunits are superposed, MQ_1_ and MQ_2_ occupy the space where hemes *b*^558^ and *b*^595^ are located in AppC subunit, respectively (Fig. 2b,c and Supplementary Fig. S5). MQ_3_ is located between h1, h2, and h9 of AppB and h4 of AppC, with the isoprenoid tail wrapped around h9 in a single loop, at the interface between the protein and the detergent micelle. The headgroup of MQ_3_ is buried and forms contacts with several non-polar residues (V61^B^, W62^B^, Y65^B^, V68^B^, I69^B^, V306^B^, Val307^B^ Ala310^B^ of AppB, and A111^C^, G114^C^, and L115^C^ of AppC). The *edge-to-edge* distance between heme *d* and MQ_3_ is 7.3 Å. Interestingly, the headgroup of MQ_3_ intersects the putative proton channel of AppB at the cytoplasmic site. Based on their proximity to the putative O_2_ pathways (see Fig. 5d), it is possible that the MQ_1_-MQ_3_ could regulate O_2_ diffusion along specific sites (see below and Supplementary Fig. S17a). In addition to the structural MQ sites, we observe a clearly resolved cardiolipin molecule at the cytoplasmic side of AppC between h5 and h9 (Fig. 2a).

**Fig. 5.**
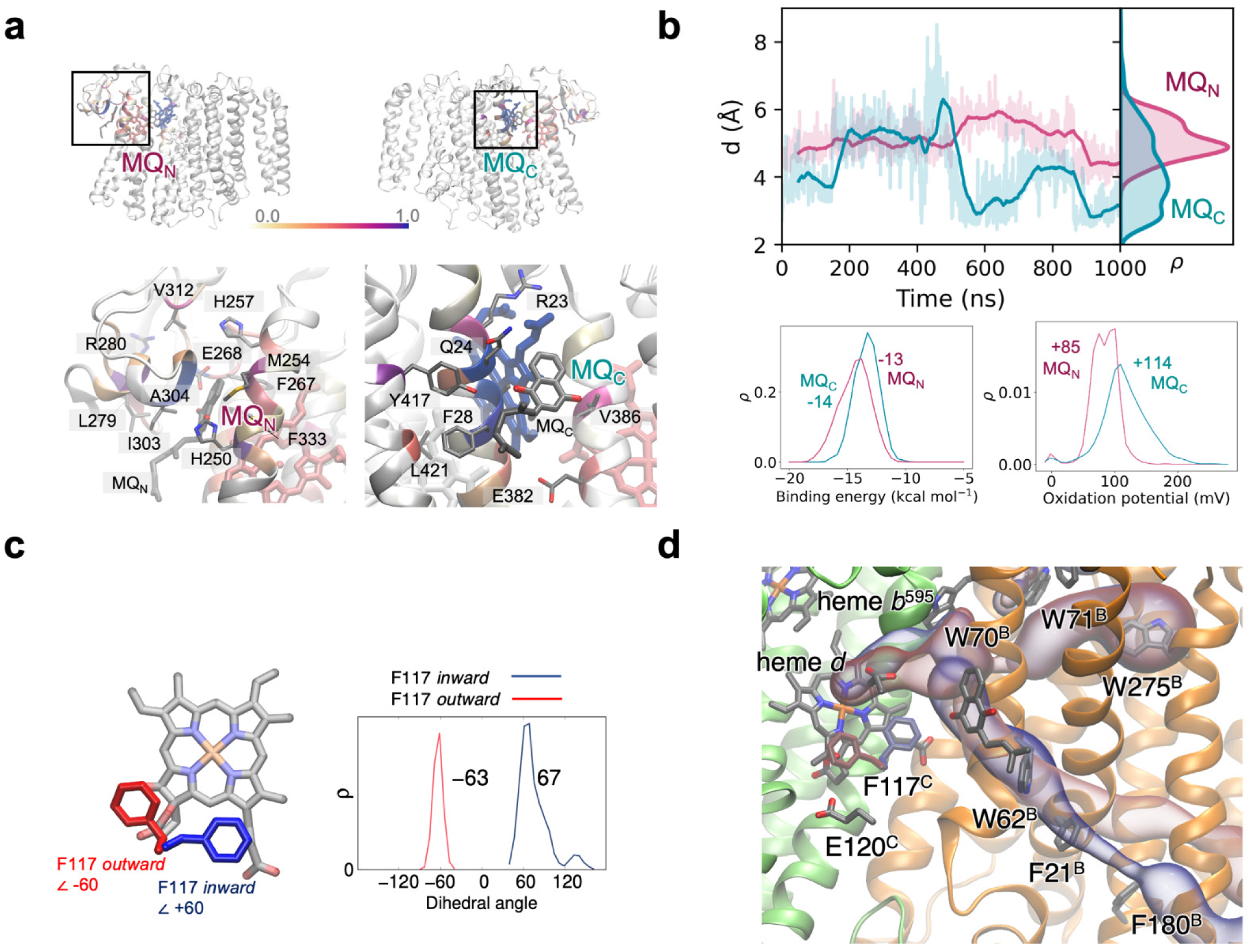
Functional analysis of *Ms* cyt *bd*-II from MD simulations. **a,** MD simulations of putative menaquinone binding sites. MQ_N_ interacts with E268^C^ and H250^C^, and surrounding non-specific residues (M254^C^, H257^C^, I303^C^, A304^C^). MQ_C_ interacts with the propionates of heme *b*^595^ and E382^C^, and with surrounding non-specific residues (Q24^C^, A27^C^, F28^C^, Y417^C^). **b,** Dynamics of modelled menaquinol (MQ) within the Q-loop. Both MQ_C_ and MQ_N_ remain close (*ca.* 4-6 Å) to heme *b*^595^, but MQ_N_ forms a more stable binding pose relative to MQ_C_. *Bottom*: calculated binding energies and oxidation potentials for MQ_N_ and MQ_C_ from MD simulations (see *Supplementary Methods*). **c,** MD sampling of the *inward* (+60°, blue) and *outward* (-60°, red) conformation of F117^C^, with histograms of the dihedral angle (N-Cα-Cβ-Cγ) based on 500 ns MD simulations of each state. **d,** Putative O_2_ pathway along the conserved residues W62^B^, F21^B^, and F180^B^. Another putative pathway forms along W70^B^, W71^B^, and W275^B^. The O_2_ tunnels are depicted in blue and red, and correspond to the *inward* and *outward* orientations of F117^C^, respectively.

### Exploration of menaquinone binding to the Q-loop

The Q-loop region (residues 259-326 between h6 and h7 of AppC) is implicated in quinol oxidation^10^. All qOR-2s, including the *Ms bd*-II, comprise a short Q-loop, that reaches out to the hydrophilic periplasm (Fig. 2, Supplementary Fig. S12). Although AppC and AppB have an otherwise nearly identical fold, the Q loop is only present in AppC. Our cryo-EM map reveals no density for the Q-loop, suggesting that the region is highly dynamic or disordered, as also observed in most other *bd* structures^10^. To obtain insight into substrate menaquinol binding, the Q-loop was explored by MD simulations based on our cryo-EM structure, in combination with an AlphaFold2^29^ model of the local unresolved parts of the structure (see *Methods*, Supplementary Table S5, Supplementary Fig. S12). Consistent with the missing cryo-EM density, the MD simulations reveal a highly dynamic Q-loop relative to the rest of the protein (Supplementary Fig. S13) that stabilizes upon menaquinone binding (Supplementary Fig. S13f). To explore the substrate binding sites, menaquinol was modelled in two initial positions: MQ_N_, based on a resolved structure of aurachin in *Ec bd*-II^22^ at *ca.* 3 Å (edge-to-edge) from heme *b*^558^, and at MQ_C_ based on a menaquinol found in *Mt bd*-I^23^ at *ca*. 3 Å (*edge-to-edge*) distance from heme *b*^595^ (Fig. 5a, Supplementary Fig. S15a,b). During the MD simulations, MQ_N_ remains at *ca.* 6 Å from heme *b*^558^, which could support rapid electron transfer (Fig. 5c). The surrounding residues, E268^C^, H250^C^, H257^C^, M254^C^, and I303^C^ interact with MQ_N_ (Fig. 5a, Supplementary Fig. S14), with the highly conserved E268^C^ (Supplementary Fig. S9) possibly functioning as a proton acceptor during the quinol oxidation process. The Q-loop motif contains several protonatable residues and water molecules, indicating that this region could support proton release to the periplasmic side of the membrane (Fig. 6a, Supplementary Fig. S5, S15, S17). MQ_C_ remains around 4-6 Å from heme *b*^558^ (Fig. 5a,b), but more dynamic relative to MQ_N_. In MQ_C_, the quinol headgroup interacts with a propionic group of heme *b*^558^ and the conserved E382^C^, which could facilitate proton transfer during the quinol oxidation. Electrostatic calculations further suggest that both sites have comparable binding energies and one-electron oxidation potential (Fig. 5b).

**Fig. 6.**
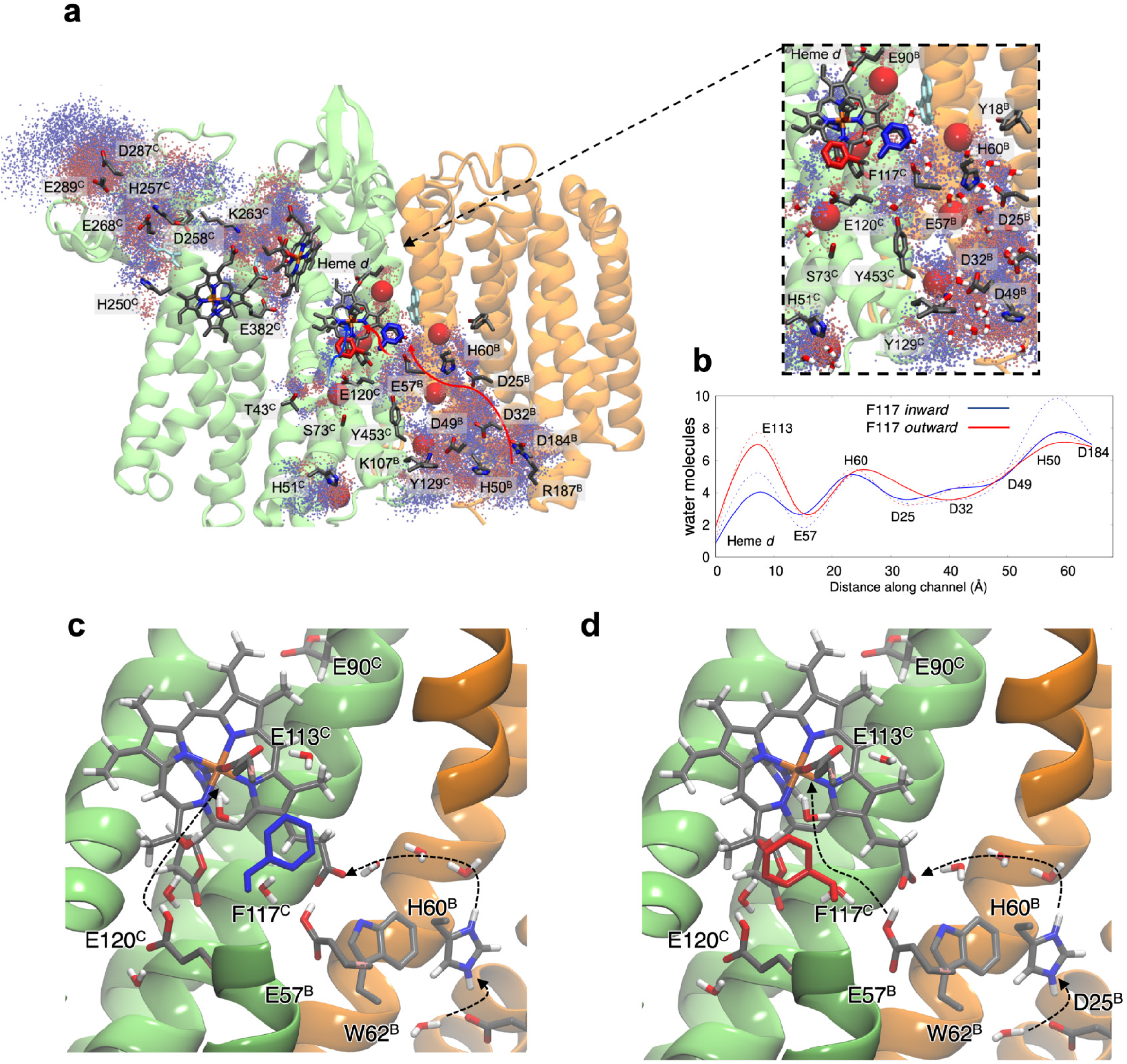
Proton pathways explored by MD simulations. **a,** Water molecules around 4 Å of the marked residues for the *inward* (blue) and *outward* (red) orientation of F117^C^. *Inset*: closeup of the heme *d* region with water clusters from MD simulations compared to water molecules resolved from cryo-EM (large red spheres). Water molecules, from MD snapshot, are shown as sticks. **b**, Average number of water molecules from MD simulations along the AppB/AppC channel, indicating that the *outward* configuration of F117^C^ enhances the hydration near E113^C^. The orientation of F117^C^ together with the protonation state of E120^C^ (dashed lines), control the hydration level around the propionate of heme *d*. **c, d,** Putative proton pathways from heme *d* based on QM/MM optimisation of the hydrogen bonded structure in the F117^C^ *inward* and *outward* conformations.

### Proton pathways and oxygen channels

To obtain insight into possible proton pathways, we next analysed the position of water molecules and protonatable residues within the resolved structure. Several water molecules (*N*=7) were resolved in the cryo-EM map, outlining two putative proton pathways (Fig. 4), which lead from the hydrophilic cytoplasmic interface region of AppB (H50^B^, D184^B^, R187^B^) and continue via Y129^C^ and K107^B^ towards heme *d*.

We next performed MD simulations to gain a more detailed insight into water dynamics and the proton pathways. The simulations reveal two distinct pathways, with *N*∼80 transient water molecules leading from the cytoplasmic side to heme *d* that align well with the water molecules resolved in the cryoEM maps (Fig. 6a). One of the pathways starts at the cytoplasmic surface of AppB, and it comprises several conserved carboxylates and polar residues (D184^B^, D32^B^, H50^B^, D25^B^, H60^B^, E57^B^) that line up along a pathway towards the heme *d* propionate and E113^C^(Fig. 6a). E113^C^, E57^B^, and D25^B^ are conserved within the complete *bd* family, while D32^B^, H60^B^, and D184^B^ are uniquely conserved within qOR-2s (Supplementary Fig. S9), thus supporting their functional role and evolutionary adaptions within the *bd* oxidases (see Discussion). The other pathway is located within AppC and connects from the surface H51^C^ via water molecules to the conserved E120^C^, leading further to heme *d* /E113^C^, although the hydrogen-bonded connectivity between E120^C^ and H51^C^ is rather weak (Fig. 6a). The water molecules observed in the MD simulations are generally highly mobile (Fig. 6a, Supplementary S17), consistent with the overall rather low water occupancy in the cryo-EM maps.

MD simulations were also carried out to explore possible O_2_ pathways leading to the heme *d* active site (see *Methods*). The O_2_ molecules distribute during the simulations near heme *b*^595^, heme *d,* and MQ_3_, with several O_2_ molecules observed near tryptophane and phenylalanine residues at the interface of AppB and AppC (Supplementary Figs. 5d, S17). The simulations support two distinct O_2_ pathways, one near the periplasm and connecting heme *b*^595^/heme *d* via a cluster of aromatic residues (W70^B^, W71^B^, W275^B^, Fig. 5b, Supplementary S17a), whereas the other pathway extends from via F117^C^, F118^C^ and W62^B^, F21^B^, F180^B^ towards the cytoplasmic side. W62^B^ is conserved within qOR-1 and qOR-2 (Supplementary Fig. S9), and previously suggested to be involved in O_2_ delivery in qOR-1s (see Ref.^21^). W62^B^ is replaced by a phenylalanine in qOR-3, indicating possible differences in the O_2_ pathways, whereas W275^B^ is unique for the qOR-2s, and support molecular adaptations within this subfamily. Taken together, our MD simulations in combination with sequence analysis suggest that the Ms *bd*-II employs O2 pathways leading to heme *d.* Interestingly, our QM/MM calculations suggest that the penta-coordinated heme *b*^595^ can also bind dioxygen without steric clashes to the surrounding protein (Supplementary Fig. S20), although our MD simulations suggest heme *b*^595^ is not connected to the O_2_ channels (Supplementary Fig. S17a).

### Mechanism of O_2_ binding and the gating role of Phe117^C^

E113^C^ coordinates the heme *d* iron in our cryo-EM structure, with a E113^C^–Fe distance of around 2.3 Å. To enable O_2_ binding and reduction, the carboxylate ligand must dissociate from heme *d*, forming a penta-coordinate Fe^2+^ with an open binding site. This was explored by hybrid quantum/classical (QM/MM) calculations to probe the principles underlying the O_2_ binding. Our QM/MM models with E113^C^ and H33^C^ ligated to the heme *d* iron (Supplementary Fig. S19) is in excellent agreement with our resolved cryo-EM density (Fig. 6, Supplementary Table S6), with the proton transfer from E120^C^ to E113^C^ favoring dissociation of the carboxylate ligand and opening of the iron coordination site (Supplementary Fig. S6, Fig. S19 Supplementary Table S6), which favors heme *d* reduction and O_2_ binding. Our MD simulations further reveal that protonation of E113 favors O_2_ binding to heme *d* (Supplementary Fig. S21). Interestingly, proton transfer to E113^C^ is possible only in the *inward* conformation of F117^C^ (Fig. 6c), whilst the *outward* conformation of F117^C^, favors the formation of a water array between E57^B^ and propionate of heme *d* (Fig. 6d). Moreover, E57^B^, implicated in proton transfer in the *E. coli bd*s^30^, connects via H60^B^ and D25^B^ to the cytoplasm side. This conformational switching could stabilise O_2_ binding to heme *d*, as also indirectly supported by the different conformations of the phenylalanine residue adopted in the qOR-1s and qOR-3s^19^ (Supplementary Fig. S8, Supplementary Table S4, see below and Discussion).

Atomistic MD simulations were performed to further probe the link between the conformational switching of F117^C^ and dissociation of the carboxylate ligand. The simulations were initiated from the two distinct F117^C^ conformations (Fig. 3), modeled both in the *apo* state of heme *d* (E113^C^ ligated), and with O_2_ bound (E113^C^ dissociated). When E113^C^ is ligated to heme *d*, F117^C^ clusters into the distinct *inward* (+60°) and *outward* (-60°) conformations, consistent with our cryo-EM data. In contrast, in the O_2_ bound form, F117^C^ becomes more dynamic and undergoes conformational switching favoring the *outward* state (Fig. 5c, Supplementary Fig. S16), which also strongly affects the hydration state of the putative proton pathway leading to the heme *d* (Fig. 6b, Supplementary Fig. S16) via the proton donors E120^C^ and E57^B^ (Fig. 6c, d) favouring the hydrogen-bonded contacts via AppB (see above). Taken together, our combined findings suggest that conformational switching of F117^C^ regulates both O_2_ and proton access to the active site, with a proton-transfer driven dissociation of the carboxylate (E113^C^) favoring the O_2_-binding to heme *d*.

## Discussion

Apart from their main role in respiratory PMF generation under lower O_2_ conditions, cyt *bd*s have been implicated in oxygen scavenging in photosynthetic and nitrogen-fixing bacteria to protect, *e.g.*, O_2_-sensitive nitrogenase^9^. In *M. smegmatis*, the *bd*-II is more upregulated upon lowering of the O_2_ concentration than the *bd*-I (Fig. 1c), similar to the role of the *bd*-II in *E. coli*, which is also upregulated at low O_2_ levels^31,32^. In general, the qOR-2 type *bd* oxidases are common in the nitrogen-fixing *Frankia* (Fig. 1b) species of Actinomycetota, consistent with a primary role as oxygen scavengers.

All cyt *bd* structures determined to date are members of the qOR-1 subfamily^16,19,20,23,24^ with the exception of one structure from the qOR-3 family (the *G. thermodenitrificans bd* ^25^), while our current study reveals a structural basis for the function of qOR-2 *bd* oxidases.

The reversal of the heme *d*/*b*^595^ positions between the *E. coli bd* (qOR-1) and the *Gt bd* (qOR-3) was suggested to originate from the insertion of an additional amino acid (L101^A^) between E99^A^ (heme *d* ligand) and E107^A^ (in *E. coli bd-I* numbering^19^), leading to a ‘bulging’ of the helix, suggested to pull E99 away from heme *d*. However, both qOR-3 and qOR-2 lack this additional residue, whilst in the Gt *bd*, the corresponding glutamate ligates to heme *b*^595^. Our structure reveals that heme *d* occupies the same space in the *Ms bd*-II as in the qOR-1 structures (Fig. 3), and shows that the additional amino acid alone does not determine the heme arrangement. The switched positions for the heme *d* was also suggested to be linked to the different conformation of the F104^A^ (*E. coli bd*-I numbering) observed between qOR-1 and qOR-3 (Supplementary Fig. S8). In the qOR-1, F104^A^ was suggested to stabilise O_2_ binding to heme *d*^19^. In this regard, our data revealed two conformations of the corresponding F117^C^, further suggesting that the residue is involved in regulating O_2_ access and binding to heme *d*.

Possible O_2_ pathways leading to the heme *d* is supported by several conserved aromatic residues present in both qOR-1 and qOR-2 subfamilies, or residues that are unique for qOR-2. Moreover, the qOR-3 subgroup has heme *b*^595^ at the position of heme *d*, further supporting that heme *d* comprises the active site in the *Ms bd*-II. However, since heme *b*^595^ is a high-spin penta-coordinated heme ligated by the conserved E389^C^, there is a long-standing question about its role in the *bd* superfamily. Both the presence of CydH, blocking direct access to the heme *b*^595^ in *E. coli bd*-I, and a nearby phenylalanine residue (F11^A^ in *Ec bd*-I, Supplementary Fig. S7) could restrict ligand access^19^. This phenylalanine is conserved within qOR-1, but exchanged for a methionine (M26^C^) in *Ms bd*-II and conserved throughout the qOR-2 subfamily, whereas it is more variable in the qOR-3 (Supplementary Fig. S9), where the heme *d* occupies this site.

The open coordination sphere and exchange of the phenylalanine, suggest that heme *b*^595^ could act as an alternative O_2_-binding site, enabled by the methionine substitution. Indeed, our QM/MM calculations suggest that heme *b*^595^ can bind dioxygen (Supplementary Fig. S20), although our MD simulations speak against a viable O_2_ channel leading to this site (Supplementary Fig. S17a), together with the lack of proton-transfer competent residues that could support the oxygen reduction chemistry.

In *E. coli bd*-I, putative proton transfer pathways from the cytoplasmic surface to the heme *d* active site consisting of hydrophilic/protonatable amino acids and water molecules were found along the CydA/B interface^10^. These pathways converge at D58^B^ and continue via the conserved E107^C^ to heme *d*, with similar pathways also found in other *cyt bd*s^16,20–23^. Functional data from site-directed mutagenesis of the *E. coli bd*s, show that E107^C^, located 7 Å from the heme *d*-Fe, is crucial for activity^33^, while recent studies also support the importance of D58^B^ and D105^B^ of CydB^30,34–36^. D58^B^ is conserved across the entire *bd* family^30,35^, whereas D105^B^ is conserved only within qOR-1s, but absent in qOR-2 and qOR-3^35^, suggesting that these *bd* subclasses employ alternative proton transfer pathways. In this regard, our cryo-EM structure and MD simulations resolved water molecules (Fig. 3) in an alternative pathway in the AppB subunit that could support proton uptake to heme *d* (Fig. 6).

Heme *d* in qOR-2 occupies the same location as in the qOR-1 (Fig. 3), but with our cryoEM data revealing a six-coordinated heme *d*, requiring E113^C^ dissociation to enable O_2_ binding. It should be noted that the optical signal for heme *d*^2+^-O_2_ at ∼650 nm is present in the air-oxidized *Ms bd*-II, but to a smaller extent than in *E. coli bd*-I (Supplementary Fig. S3). Our simulations further suggest that the conformation of F117^C^ regulates O_2_ and proton access to heme *d* (Fig. 5 and Supplementary Figs. S16, Fig. S17c). In this regard, we suggest that reduction of heme *d* couples to protonation of E113^C^, which favors dissociation of the glutamate ligand and induces O_2_ binding, which in turn flips F117^C^ to the *inward* position (Supplementary Fig. 21). The suggested conformational gate is expected to favor O_2_ binding at low pH, where E113^C^ is protonated. The conformational changes could help to kinetically trap O_2_, which is particularly important at microaerobic environments (*cf*. also Refs. ^37,38^).

Ligand switching via the suggested carboxylate shift mechanism allows O_2_-binding by controlling open metal coordination positions. Similar mechanisms are commonly observed in di-metal carboxylate proteins, in various model complexes^39,40^, as well as in certain β-carbonic anhydrases, where the dissociation of a Zn^2+^-bound glutamate is induced by pH changes^41^. Ligands switching also occurs in some heme peroxidases, where a histidine ligand dissociates during catalysis^42^, thus indirectly supporting the conformational switching enabling O_2_ binding in qOR-2.

Taken together, our findings reveal a molecular basis for the function of the *bd*-II oxidase of *M. smegmatis* and elucidate the location of proton and dioxygen pathways and molecular gates near the active site that could support a functional role of qOR-2 type *bd* oxidases as efficient oxygen scavengers in micro-aerobic environments.

## Data availability

The cryo-EM map and the atomic model for the *Ms bd*-II were deposited to the EMDB and PDB, respectively, with the following accession codes: EMD-53529, and 9R2G. MD data is deposited at zenodo: 10.5281/zenodo.15385662.

## Acknowledgements

This work was supported by the Knut and Alice Wallenberg Foundation (KAW 2019.0043 to VRIK, MH, and PÄ, KAW 2023.0201 to MH and KAW 2019.0251, 2024.0220 to VRIK), the Swedish Research Council (2020-04081 to V.R.I.K, 2019-04124 to PÄ and 2021-03992 to MH), and the Göran Gustafsson Foundation (to VRIK). The authors are thankful for the computing time provided by the National Academic Infrastructure for Supercomputing in Sweden (NAISS 2024/1-28; 2023/1-31). The cryo-EM data were collected at the Cryo-EM Swedish National Facility funded by the Knut and Alice Wallenberg, Family Erling Persson and Kempe Foundations, SciLifeLab, Stockholm University and Umeå University.

## Author contributions

Design of study: MJ, TK, APGH, DS, VRIK, MH, and PÄ; Data acquisition: MJ, TK, APGH, SS, DL, JV; Data analysis: MJ, TK, APGH, SS, DL, DS, JV, VRIK, MH, and PÄ; Writing, original draft: MJ, TK, APGH, and PÄ; Writing, review and editing: MJ, TK, APGH, SS, DL, DS, JV, VRIK, MH, and PÄ; Funding acquisition: VRIK, MH, and PÄ.

## Abbreviations

HCuOs: heme-copper oxidases
qOR1-4a: quinol/oxygen *bd* oxidoreductases 1-4a
LMNG: lauryl maltose neopentyl glycol
cryo-EM: cryogenic electron microscopy
F117^C^: phenylalanine 117 in AppC subunit
*Ms*: *Mycobacterium smegmatis*
*Gt*: *Geobacillus thermodenitrificans*
*Ec*: *Escherichia coli*
*bd*: *cytochrome bd oxidase*
Q-loop: quinol binding loop
Q: quinol
MQ: menaquinol
h: helix
MD: molecular dynamics
QM/MM: quantum mechanics/molecular mechanics.

## Methods

### Bioinformatic analysis

All Genome Taxonomy Database GTDB (R09-RS220; ^43^) species representative genomes were ORF-called with Prokka (v.1.14.6^44^). Subsequently, the genomes were searched with the HMMER (v.3.4^45^) HMM profiles for the different types of AppC/CydA as reported by Murali *et al*. at <u>GitHub</u> together with Ref.^3^). For every protein found, the best scoring profile was selected as its annotation. A minimum score of 300 was applied to the hits. To investigate amino acid substitution patterns at important positions of the *Mycobacterium smegmatis* AppC and AppB proteins, all CydA proteins matching the profiles from Ref ^3^ were collected (*n*=54743), clustered with Usearch (v.11.0.667^46^) at an identity threshold of 0.6 (*n*=3022) to reduce the degree of redundancy and aligned with Clustal Omega (v1.2.4; ^47^). Since there is no corresponding classification of the AppB/CydB proteins, we classified potential proteins by first identifying them with the Pfam^48^) profile PF01654 using HMMER. AppB/CydB proteins encoded by genes located within a distance of five genes from a corresponding AppC/CydA-coding gene were classified in correspondence with its nearby CydA class. This resulted in 2007 proteins that were aligned with Clustal Omega. Interesting positions in both alignments were then extracted and converted to amino acid proportions. Figures were produced with R (v.4.3.3^49^) using the Tidyverse packages (2.0.0^50^).

### Quantitative real-time reverse transcriptase PCR

All the chemicals, enzymes, RT-qPCR reagents, SYBR RT-qPCR master mix and cDNA synthesis kit were purchased from Thermo-Fisher Scientific, USA. The DNA oligonucleotides were synthesized by Eurofins genomics. The *M. smegmatis* mc^2^155 strain was grown in liquid 7H9 Middlebrook broth (Merck, USA), supplemented with 0.05% Tween 80, and 10% albumin-dextrose-NaCl (ADS) (henceforth referred to as 7H9 broth), while the cells were grown on 7H10 agar (4.7 g L^-1^) plates supplemented with 10% ADS. The starter cultures for RNA isolation were prepared by inoculating a single colony of *M. smegmatis* mc^2^155 into 15 mL of 7H9 broth and grown at 37°C in a shaking incubator (220 rpm), until the cultures reached an OD_600_ of 1. Then 1.5 mL of the cell suspension was inoculated into 30 mL of 7H9 Middlebrook medium to give an OD_600_ of 0.05. For the slow-growing cells, culture flask lids were tightly screwed and gently stirred at 170 rpm while for the fast-growing cells, culture flask lids were loosely screwed and stirred at 250 rpm. The cultures were harvested at the time points corresponding to log phase. The expression of *appC, appB, cydA* and *cydB* in *M. smegmatis* mc^2^155 cells, grown under slow or fast conditions, were determined by quantitative real-time PCR using specific primers (Supplementary Table S1). The total RNA was isolated from the cells using the TRIzol Max Bacterial RNA Isolation Kit as per the manufacturer’s instructions. The quantity and quality of RNA was checked using a nanodrop and then running an RNA gel. The RNA preparations were further incubated with DNase I (1 U µg^-1^) to remove traces of DNA and then used for cDNA synthesis by utilizing the RevertAid First Strand cDNA Synthesis Kit according to the manufacturer’s instructions. Semi-quantitative PCR reactions were performed using the Taq DNA polymerase and 15-20 pmol gene-specific primers. The cycling program comprised initial denaturation at 94°C for 3 min, denaturation at 94°C for 15 sec, primer annealing at 60°C for 30 sec and extension at 72°C for 30 s. The RNA samples without reverse transcriptase were included as a negative control. Real-time PCR was done in 15 μL volume using 1X SYBR Green Master Mix (Applied Biosystems), 10 pmole of gene specific primers and 1μL of cDNA. The reaction was performed in STEP-ONE System (50°C, 2 min; 95°C, 10 min, 1 cycle; 95°C, 15 s; 60°C 1 min, 40 cycles). Following amplification, a melting curve analysis was performed to verify the authenticity of the amplified product. SigA quantification was used as an internal control for normalization. Fold differences of mRNA levels over control values were calculated.

### Plasmid construction and bacterial cell cultures

Open reading frames MSMEG_5605-5606 (*appCB*) of M .smegmatis mc2155 were amplified using PCR primers: AGTACATATGGTGTTCACCGAAACCCTGCTTC and AGCAAAGCTTGCGTGACCACTCCTCGGTCTGC) and cloned in the NdeI/HindIII digestion sites of a modified pST-K vector^51^ (N-terminal His-tag removed, C-terminal Twin-Strep-tag added); yielding the pST-K-appCB_Twin-Strep_. DNA sequencing was performed (Eurofins) to confirm the cloning. The wild-type strain MC^2^155 of *M. smegmatis* was transformed via electroporation with the plasmid pST-K-appCB_Twin-Strep_, with the following settings: 2.5 kV, 1000 W, 25 mF. Cultures of *M. smegmatis* were cultivated in 7H9 medium (Difco), enriched with TDS (10 g L^-1^ tryptone, 2 g L^-1^ dextrose, 0.8 g L^-1^ NaCl). A preculture of 15 mL was initiated by inoculating with cells previously grown on an agar plate and was subsequently incubated at 30°C with continuous shaking at 180 RPM for 24 h. This preculture served as the inoculum for a larger culture (2 L in a 2.5 L full-baffled flask-Tunair™), which was further incubated semi-aerobically (achieved by lowering the shaking speed to 100 RPM) for an additional 48 h. The cells were harvested by centrifugation at 5900 x g for 25 min (rotor J.LA 8.1) and flash-frozen using liquid nitrogen for storage. The cell pellets were kept at -20°C until further experimentation.

### Purification of the *M. smegmatis* cytochrome *bd*-II oxidase

All purification processes were carried out at 4°C. Approximately 50 g of wet cell mass was homogenized in 300 mL of buffer A (50 mM HEPES, pH 7.4, 150 mM NaCl), supplemented with one tablet of cOmplete™ Mini Protease Inhibitor Cocktail (Roche) and a small amount of DNase I. Cell disruption was achieved through three to four passes at 35 kPsi in a cell disruptor (Constant Systems Ltd.). Following disruption, the mixture was centrifuged to spin down cell debris at 15,500 x g for 15 min. The cell membranes were separated by ultracentrifugation at 215,000 x g for 90 min. The recovered membranes (5-10 g) were then resuspended in buffer A to a final protein concentration of 10 mg mL^-1^ (determined by Lowry assay^52^ and solubilized by the addition of 1% (w/v) lauryl maltose neopentyl glycol (LMNG) while gently stirring for 60 min. Any uncrushed material was removed via centrifugation at 215,000 x g for 60 min. The supernatant was collected and subjected to affinity chromatography. The solubilized protein in the supernatant was applied to a gravity column (2 mL Strep-tactin Superflow Plus, VWR) that had been pre-equilibrated with buffer B (50 mM HEPES, pH 7.4, 150 mM NaCl, 0.003% LMNG). Once the solubilized material was loaded entirely, the column was washed with 10 mL of buffer B, followed by elution with 10 mL of buffer B containing 10 mM desthiobiotin. The eluted protein was concentrated using a 100-kDa molecular weight cutoff concentrator (Merck Millipore), flash frozen with liquid N_2_ or subjected to further experiments. The affinity-purified cyt *bd*-II was subjected to a size exclusion chromatography to ensure its homogeneity. The Superdex 200 10/300 column (Cytiva) was equilibrated with two column volumes (CV) of buffer B. Previously collected protein sample was applied on the column. Peak fractions were collected and analyzed in further experiments. The cyt *bd*-I from *E. coli* used for spectral comparisons was prepared as in Ref.^35^ based on the plasmid from Ref. ^53^.

### Optical UV-Vis spectroscopy

Optical spectra of the (air) oxidized and (sodium dithionite) reduced cyt *bd*-II oxidase were measured using Cary 3500 Compact UV-Vis spectrophotometer (Agilent). The concentration of the enzyme was determined using an extinction coefficient of 27 mM^-1^cm^-1^ for the 629 to 650 nm wavelength pair^11^.

### Activity Assays

The oxidoreductase activity of the cyt *bd*-II was assessed in a solution comprising 50 mM HEPES (pH 7.4), 150 mM NaCl, and 0.003% LMNG at 25°C, using a Clark-type electrode (Hansatech instruments). Reduced 2,3-dimethyl-[1,4]-naphthoquinone (DMNQ) served as the substrate. Background oxidation was recorded by adding 2.5-15 µl of a 20 mM stock solution of reduced quinol over a period of approximately 45 s. Subsequently, the reaction was initiated by introducing cyt *bd*-II (125 nM), and the changes in oxygen concentration were monitored for 1-2 min. The activity was calculated from the initial slope of the oxygen concentration versus time. Control measurements without the addition of cyt *bd*-II were performed to determine the background oxygen reduction rate, which was subtracted from the rate values obtained in the presence of cyt *bd*-II. DMNQ was dissolved to 20 mM in ethanol and reduced using sodium borohydride as in Ref.^54^. The reduced naphthoquinol was flash frozen in liquid N_2_ and stored at -80°C until further use.

### Mass spectrometry

The mass spectrometry protein identification analysis was prepared by Alphalyse™ using in-solution protein digestion and nanoLC-MSMS. The digestion was done on a pre-reduced and alkylated sample using trypsin. Resulting peptides were concentrated and a control using bovine serum albumin (BSA) was conducted alongside the sample. The peptides were treated with 0.1% formic acid, injected to a mass spectrometer and left for a 50-min-long gradient run. Results were run against a collective UniProtKB database giving the entries for two cyt. *bd*-II subunits (AppC and AppB) and an additional peptide with a low MS score.

### Sample preparation and cryo-EM data collection

Protein purified in LMNG (40 μM) was used for the sample preparation. The grids (Au300 mesh R1.2/1.3 grid, Quantifoil, Micro Tools GmbH, Germany) were glow-discharged in air at 20 mA for 120 s (PELCO easiGlow). 2 μL of the protein sample was blotted for 3 s at 4°C, 100 % humidity and plunge-frozen in liquid ethane using Vitrobot Mark VI (Thermo Fisher Scientific). Titan Krios G3i electron microscope (Thermo Fisher Scientific) Ceta-D camera and a Gatan K3 BioQuantum detector was used for data collection. The dataset was acquired in electron-counting mode at a nominal magnification of 130,000 (0.650 Å/pixel) and contained 17000 exposures (40 exposure fractions each). EPU software package (v2.12.1; Thermo Fisher Scientific) was used for automated data collection with camera exposure rate was 18.4 e^-^/pixel/s and total exposure of the sample 93.5 e^-^/Å^2^ (Supporting information, Supplementary Figure S1 and Supplementary Table S1).

### Cryo-EM data processing, structure building, and refinement

The dataset was processed using CryoSPARC v4^55^ with motion correction and CTF estimation performed using the Patch motion correction and Patch CTF estimation, as implemented in cryoSPARC. Initial particles used for generation of initial 2D classes, later used by the Template picker job, was picked from 1000 images using the Blob picker job in cryoSPARC. The initial set of particles (7,907,011) was reduced by 2D classification to 3,231,874 particles, followed by tree more rounds of 2D classification and *ab initio* reconstruction and one more 2D classifications to final set of 200,084 particles. Final set of particles was reextracted using the box size 360 px. Further refinement was performed using additional ab-initio job and non-uniform refinement, followed by local motion correction, second non-uniform refinement and local CTF refinement without symmetry enforced. The last non-uniform refinement resolved final map to overall resolution of 2.8 Å. The initial structure model used for map interpretation was downloaded from AlphaFold^29^ with the AppC subunit: AF-A0A653F9S4-F1-model_v4, AppB subunit: AF-A0A8B4QQ04-F1-model_v4. The model was refined against the map using Coot^56^ and real-space refinement in Phenix^57^. The figures were prepared with PyMOL (The PyMOL Molecular Graphics System, *Schrödinger LCC* (2002)).

### Molecular dynamics simulations

Atomistic molecular dynamics simulations of the *M. smegmatis bd*-II oxidase were performed based on our resolved cryo-EM structure (PDB ID: 9R2G), and by modeling the Q-loop and other missing loops using Alphafold2^29^ (uniprot code: A0A653F9S4) (Supplementary Fig. S12). A MQ molecule, observed in *Mt bd*-I oxidase (PDB ID ID: 7nkz)^23^ was modeled in the MQ_C_ site, whereas MQ_N_ was modelled based on aurachin in the *E. coli bd*-II (PDB ID:7ose)^22^. The MD simulations were performed in the *apo* state (as resolved in cryo-EM), MQ_N_/MQ_C_ models with either O_2_-bound heme *d* or E113^C^ ligated to heme *d*, as well as in the two distinct conformations of F117^C^, observed in the cryo-EM data (Supplementary Fig. S12, Supplementary Table S5). The MD models were embedded in a POPC/POPE/PI/cardiolipin lipid membrane using CHARMM-GUI^58^, solvated with TIP3P water molecules, and NaCl ions at 0.15 mM concentration (Supplementary Fig. S12), with 55 dioxygen molecules added into the simulation box (see Supplementary Methods). The cardiolipin molecule was modeled based on the cryo-EM maps (Supplementary Fig. S12), and menaquinol was modelled in the Q-loop (see above), whereas menaquinone was modeled in sites MQ_1_, MQ_2_, and MQ_3_ based on our cryo-EM data. The protein, lipids, and water molecules were described using the CHARMM36 force field^59^, whereas force fields parameters were derived for heme *b*^558^, heme *b*^595^, and heme *d* in their oxidized (Fe^III^) and reduced (Fe^II^) states and with O_2_-bound (Fe^II^-O_2_), based on quantum chemical calculations (B3LYP/def2-SVP/def2-TZVP level) (data deposited in kailalab gui^28^). Protonation states were modeled based on electrostatic calculations by computation of the electrostatic potential by the linearized Poisson-Boltzmann equation model in combination with Monte Carlo sampling (PBE/MC) of protonation states^60^ (Supplementary Table S5). The MD system comprised *ca*. 282,000 atoms (Supplementary Fig. S12), and was propagated in two replicas for 0.5 μs using a 2 fs integration step at *T*=310 K, *P*=1 bar in an *NPT* ensemble. The protein was initially fixed and gradually relaxed for 10 ns with harmonic restraints of 2 kcal mol^-1^ Å^-2^ (applied on all protein atoms and cofactor heavy atoms), followed by 1 ns simulation with harmonic restraints (2 kcal mol^-1^ Å^-2^ on the protein backbone), and 10 ns equilibration with weak restraints (0.5 kcal mol^-1^ Å^-2^) on all C_α_ atoms, followed by the unbiased MD simulations. Binding affinities for the MQ ligands (modeling MQ-1) were estimated based on a molecular mechanics/Poisson-Boltzmann surface area (MM/PBSA) continuum solvation model^61^, as implemented in APBS^62^ without considering entropic effects^63^. MQ oxidation potentials were calculated based on the MC/PBE methodology^64^, based on quantum chemical models and parametrisation (see Supplementary Methods). The MD simulations were propagated using NAMD3^65^ and the trajectories were analyzed using VMD^66^, PYMOL and CAVER^67^. See Supplementary Table S5 for list of MD simulations and modeled ligand/ protonation states.

### QM/MM simulations

QM/MM models were constructed based on the cryo-EM map (PDB ID: 9R2G) to probe possible conformations and ligand binding in heme *d*. The QM region was modelled at the B3LYP-D3/Fe def2-TZVP/def2-SVP level and coupled to the MM system (modelled at the CHARMM36) by an additive electrostatic embedding, with link atoms introduced between Cα and Cβ atoms. The QM/MM models, comprising 156 QM atoms/12,000 MM atoms, constructed by truncating the full system, were structure optimized for O_2_-bound heme *d* (Fe^II^-O_2_) or with E113^C^ ligated to Fe^III^ in *inward*/*outward* conformations of F117^C^. The models included the resolved cryo-EM water molecules and water molecules from the MD simulations. The QM/MM calculations were performed using CHARMM/TURBOMOLE^68–70^.

