## Supplementary Information for "The *Mycobacterium smegmatis bd*-II terminal oxidase employs a carboxylate shift mechanism"

**Content**

**Extended Methods**

**Figure S1**. Combination of *bd* families in species in the nine largest Actinomycetota orders.
**Figure S2**. Size-exclusion chromatography of purified *Ms* cyt *bd*-II in LMNG.
**Figure S3**. Spectral analysis of Ms *bd*-II.
**Figure S4**. Cryo-EM data of Ms *bd*-II.
**Figure S5**. Structural analysis of *bd*-II.
**Figure S6**. Cryo-EM electron density map (σ=2 level).
**Figure S7**. Ligands and environment of heme $b^{595}$.
**Figure S8**. Ligands and environment of heme *d*.
**Figure S9**. Amino acid distribution at selected positions of the *M. smegmatis* AppC (A) and AppB (B).
**Figure S10**. Comparison of Ms *bd*-II with previous *bd* structures.
**Figure S11**. Ligand binding sites.
**Figure S12**. MD setup of *bd*-II.
**Figure S13**. MD analysis of *Ms bd*-II with and without MQ in the Q-loop.
**Figure S14**. MQ interactions in the Q-loop.
**Figure S15**. Overview of titratable residues in the Q-loop.
**Figure S16**. Dynamics of $F117^C$ and hydration level from MD simulations in different ligand and protonation states.
**Figure S17**. Overview of hydration and $O_2$ accessibility from MD simulations.
**Figure S18**. Putative proton pathways from MD simulations.
**Figure S19**. Comparison of QM/MM models of heme *d* with different ligands and oxidation state with cryo-EM data.
**Figure S20**. QM/MM model of heme $b^{595}$ with bound $O_2$.
**Figure S21**. Diffusion of $O_2$ near heme *d*.

**Table S1.** *Actinomycetota species* with only qOR-2 *bd*.
**Table S2.** Data collection and processing of cryo-EM data.
**Table S3.** List of primers used for quantitative real-time PCR.
**Table S4.** List of corresponding amino acid numbers in selected *cyt* bds.
**Table S5.** List of molecular dynamics (MD) simulations.
**Table S6**. Heme *d* ligands and bond distances from QM/MM models.

### Extended Methods

#### MD simulations of $O_2$ channels

The $O_2$ distribution in *bd*-II were analyzed based on the combined 2 μs MD simulation data (Table S5, simulations S1/S2). To this end, MD simulations were initiated with a dioxygen molecule ligated to the heme *d*-Fe or next to the heme *d* in the heme *d*-E113$^C$ ligated state. Initial $O_2$ positions in the protein were obtained from steered molecular dynamics simulations, where 55 $O_2$ molecule were radially pulled from heme *d*-Fe towards tryptophane residues within the *bd*-II (W56$^B$, W70$^B$, W71$^B$, W157$^B$, W250$^B$, W275$^B$, W351$^C$, W330$^B$, W69$^C$, W95$^C$, W131$^C$, W163$^C$, W183$^C$, W292$^C$, W323$^C$, W355$^C$, W365$^C$, W395$^C$, W398$^C$, W414$^C$, W415$^C$, W442$^C$, W62$^B$) using a harmonic restrain of 10 kcal mol$^{-1}$ Å$^{-2}$ followed by a short (2 ps) MD relaxation in $T$=310 K without restraints, and the unbiased MD simulations (Table S5). The dioxygen was modeled using the force field parameters developed in Ref.[1].

#### Estimation of MQ oxidation potentials and p$K_a$ values in bd-II

Oxidation potentials ($E_m$) and p$K_a$ values were estimated based on the continuum electrostatic calculations[2]. To this end, the protein was modeled as heterogenous dielectric medium (ε=10), the surrounding water was described as a homogeneous dielectric media (ε=80), the lipid membrane, the protein and cofactors were modeled explicitly using the CHARMM36[3] force field in combination with developed force field parameters of the cofactors. In this regard, RESP charges of heme $b^{595}$, heme $b^{558}$ and heme *d* with different ligands and as well, charges of MQ states Q, $QH_2$, $QH^-$ and SQH• were calculated at the B3LYP-D3/def2-TZVP level in TURBOMOLE[4]. The $E_m$ and p$K_a$ values were calculated by solving the linearized Poisson-Boltzmann equation (PBE) using APBS in combination with Monte Carlo sampling in Karlsberg+[5,6] based on 1000 frames extracted from 500 ns MD simulation (simulation S1, Table S5).

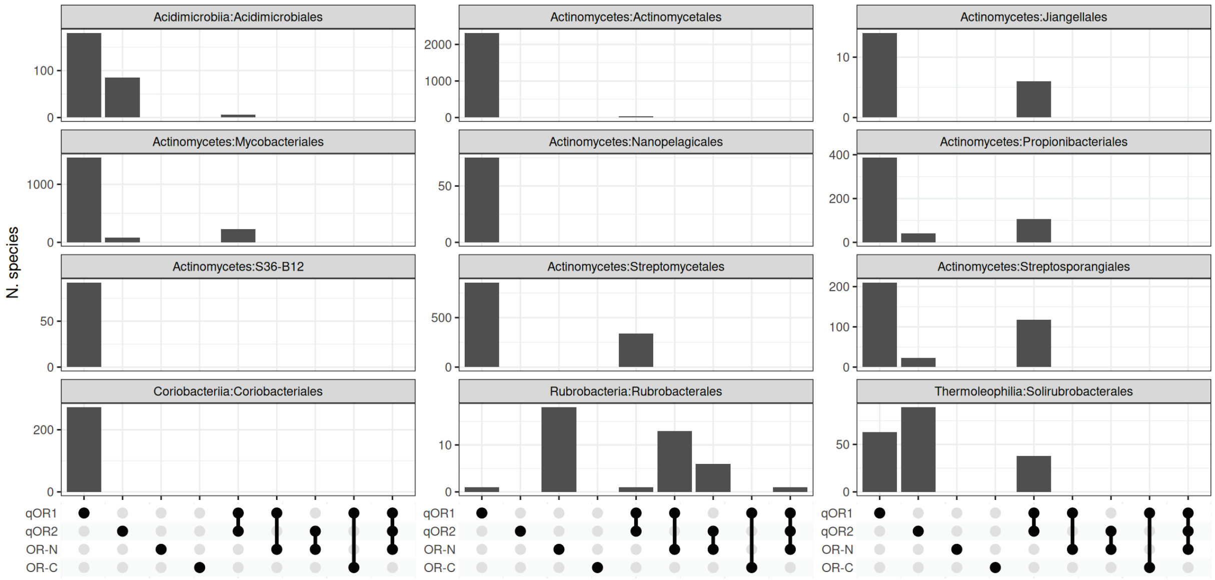


**Figure S1.** Combinations of *bd* families in species in the nine largest *Actinomycetota* orders.

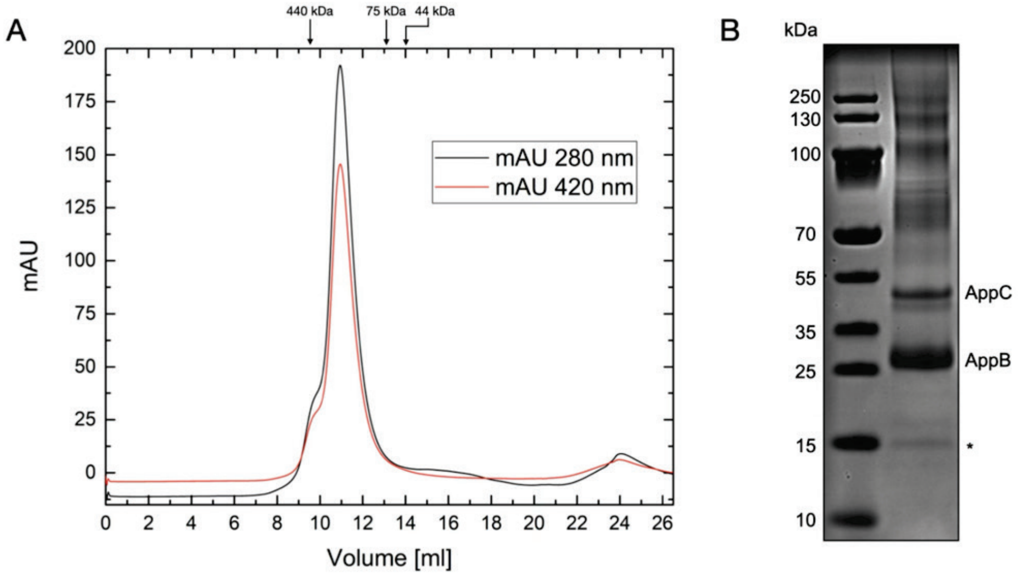


**Figure S2.** A) Size-exclusion chromatography of purified *Ms* cyt *bd*-II in LMNG. The elution peak corresponds to ~240 kDa. B) SDS-PAGE analysis of the SEC peak fraction on 4-16% gradient gel. The asterisk represents a reocurring peptide band, not assigned to any density in the structure.

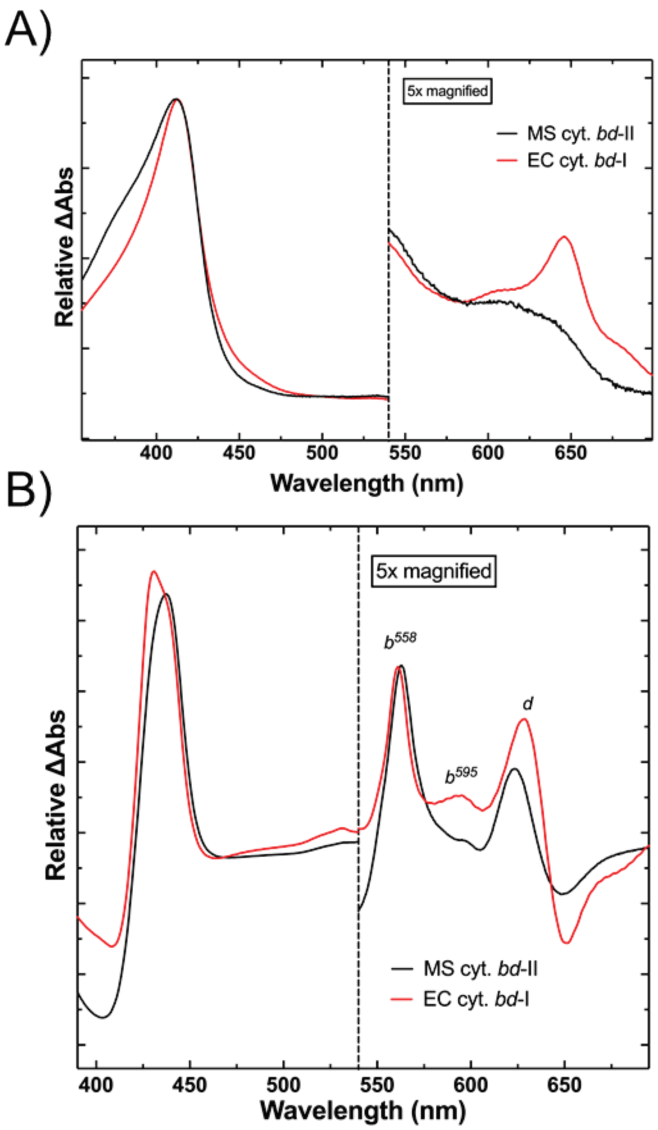


**Figure S3**. Spectral analysis of *Ms bd*-II. A) Comparison of air-oxidized heme spectra of *M. smegmatis* cyt *bd*-II (black) and *E. coli* cyt *bd*-I (red). Oxidized heme *d* absorption maxima: 650 nm for *Ec bd*-I, 646 nm for *Ms bd*-II. B) Comparison of the reduced-oxidized spectrum *M. smegmatis* cyt *bd*-II (black) and *E. coli* cyt *bd*-I (red). Reduced heme $b^{558}$ peak position: 561 nm for *Ec bd*-I, 563 nm for *Ms bd*-II. Reduced peak position for $b^{595}$: 595 nm for *Ec bd*-I, 598 nm for *Ms bd*-II. Reduced peak positions for *d*: 629 nm for *Ec bd*-I, 624 nm for *Ms bd*-II. Oxidized traces normalized to the 410 nm value in Soret region and reduced-oxidized traces normalized to the 435 nm value in Soret region. Measurements were done in 50 mM HEPES pH 7.4, 150 mM NaCl, 0.003% LMNG (w/v).

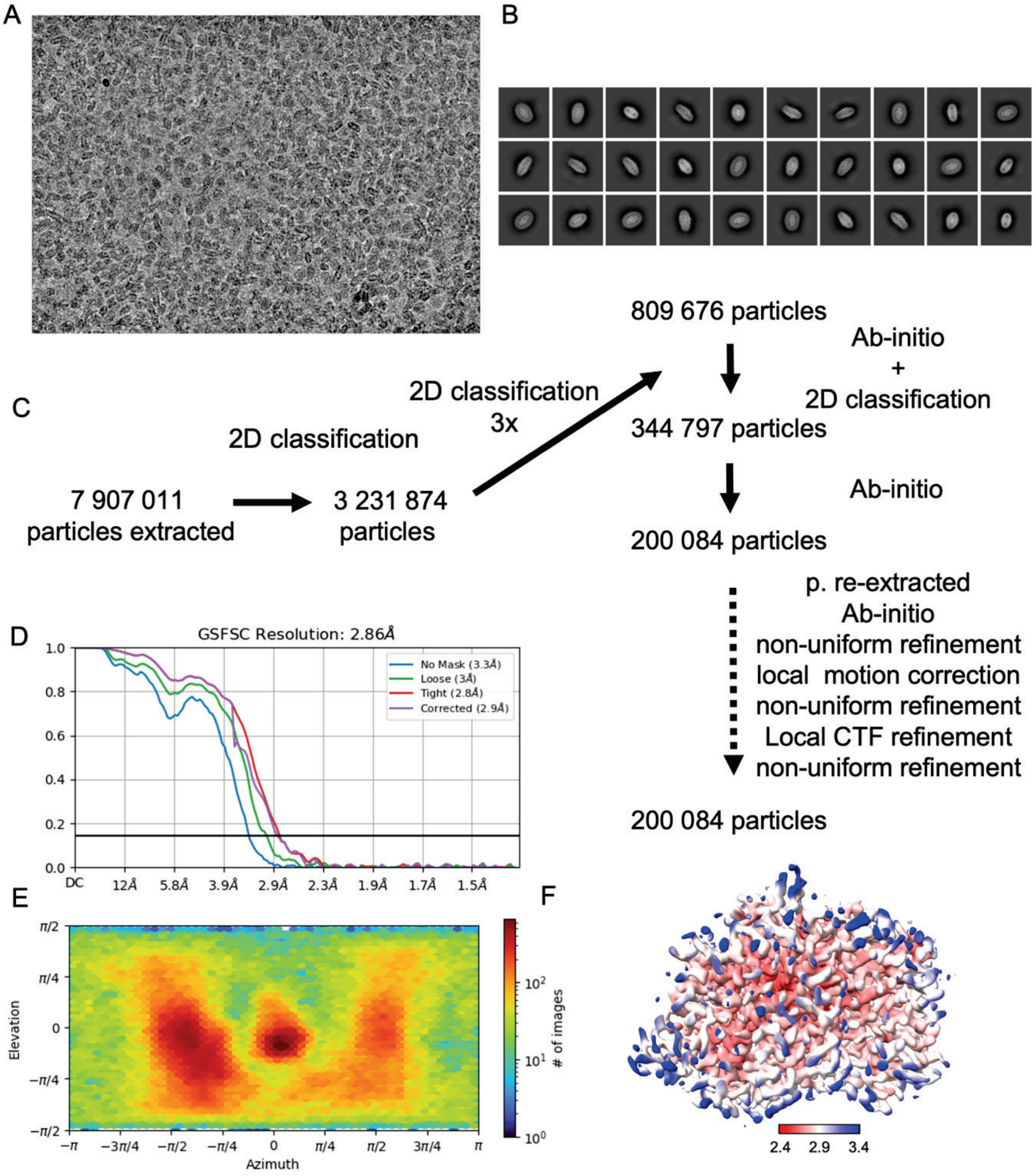


**Figure S4. Cryo-EM data of *Ms bd*-II. A)** Micrograph. **B)** 2D classes. **C)** Schematic workflow of data processing. **D)** Fourier shell correlation plot of the final 3D-refinement of particle. **E)** Distribution of viewing direction for final refinement of particles. **F)** Local resolution distribution over the final volume (from 2.4 to 3.4 Å).

A

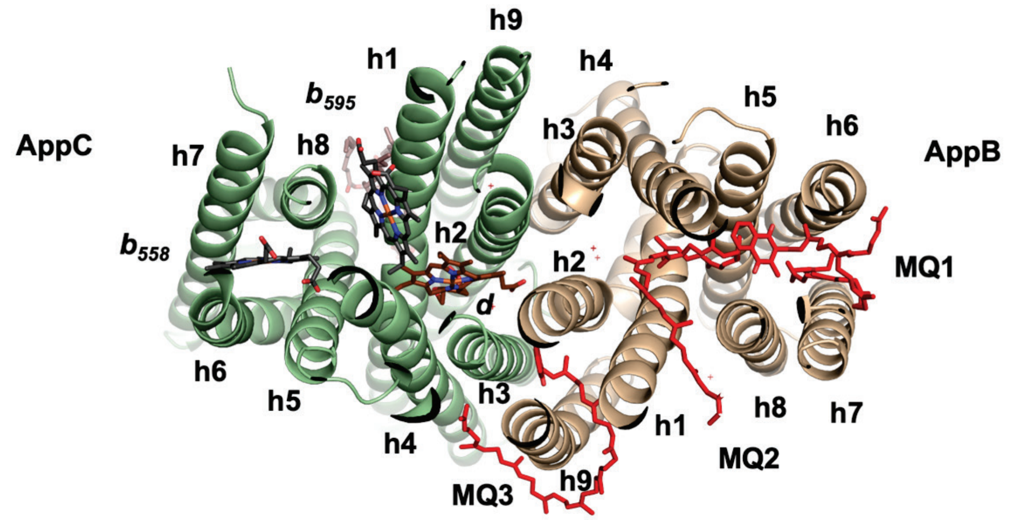


B

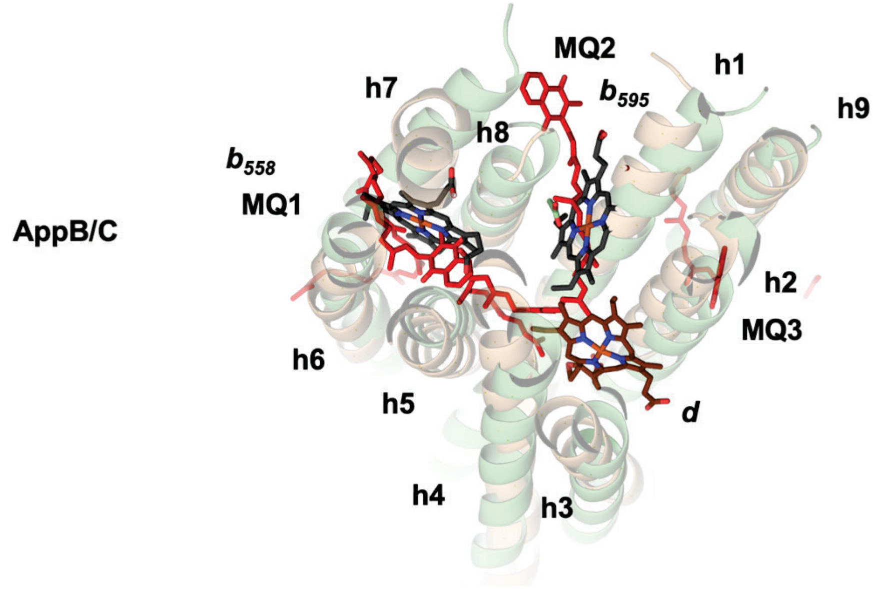


**Figure S5.** Structural analysis of *Ms bd*-II. **A)** Top view of the complex with individual helixes numbered and position of the ligands marked. B) Superposition of the AppC and AppB subunit demonstrating fold conservation. Both subunits probably evolved from the same gene. Equivalents of heme binding site in AppC are occupied by MQs (in red) in AppB.

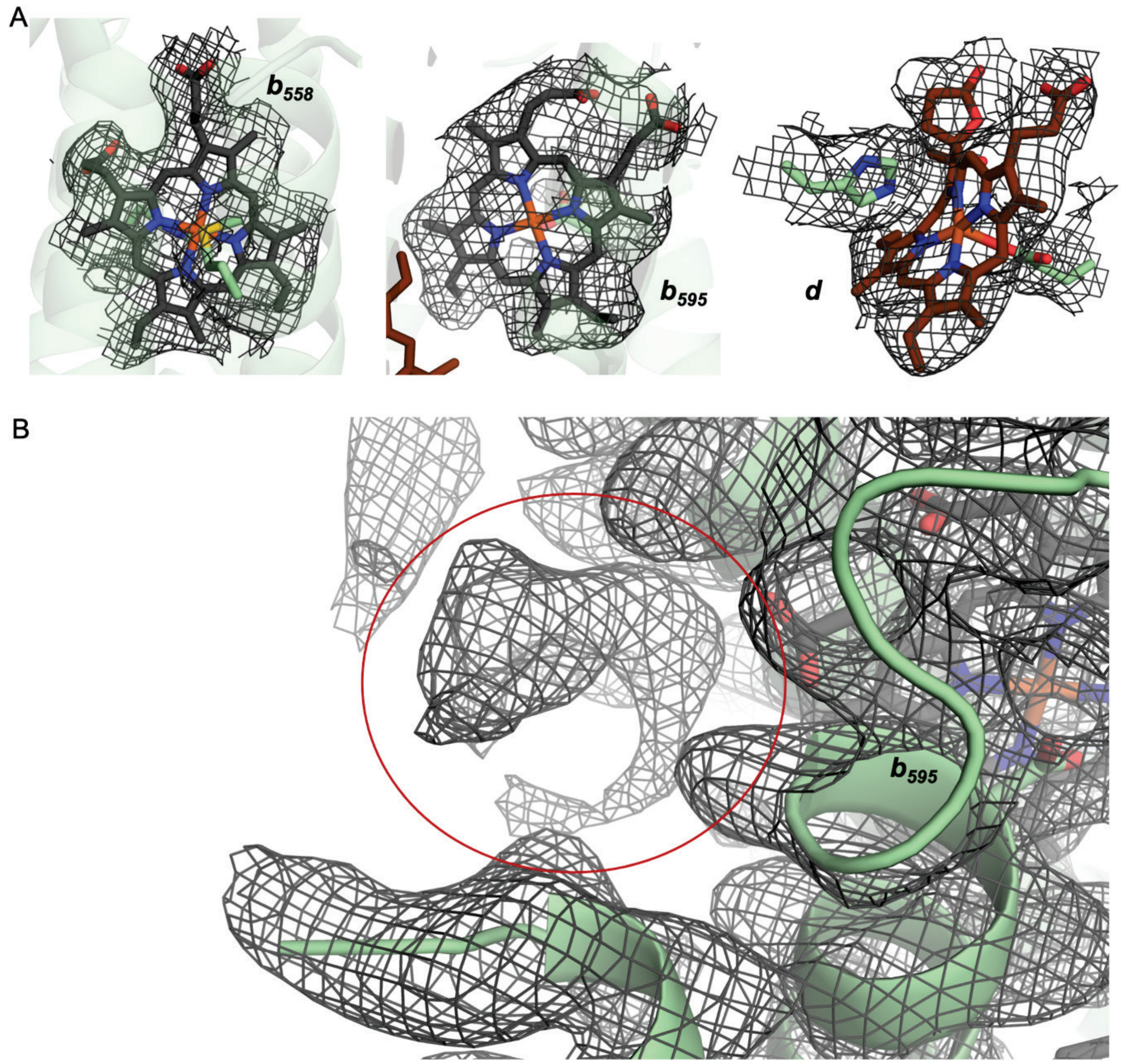


**Figure S6.** Cryo-EM electron density map (σ=2 level) of **A)** the individual hemes **B)** the unmodeled density next to heme $b^{595}$ is observed at the MQ binding site of *M. tuberculosis bd*-I (PDB ID: 7NKZ[7]).

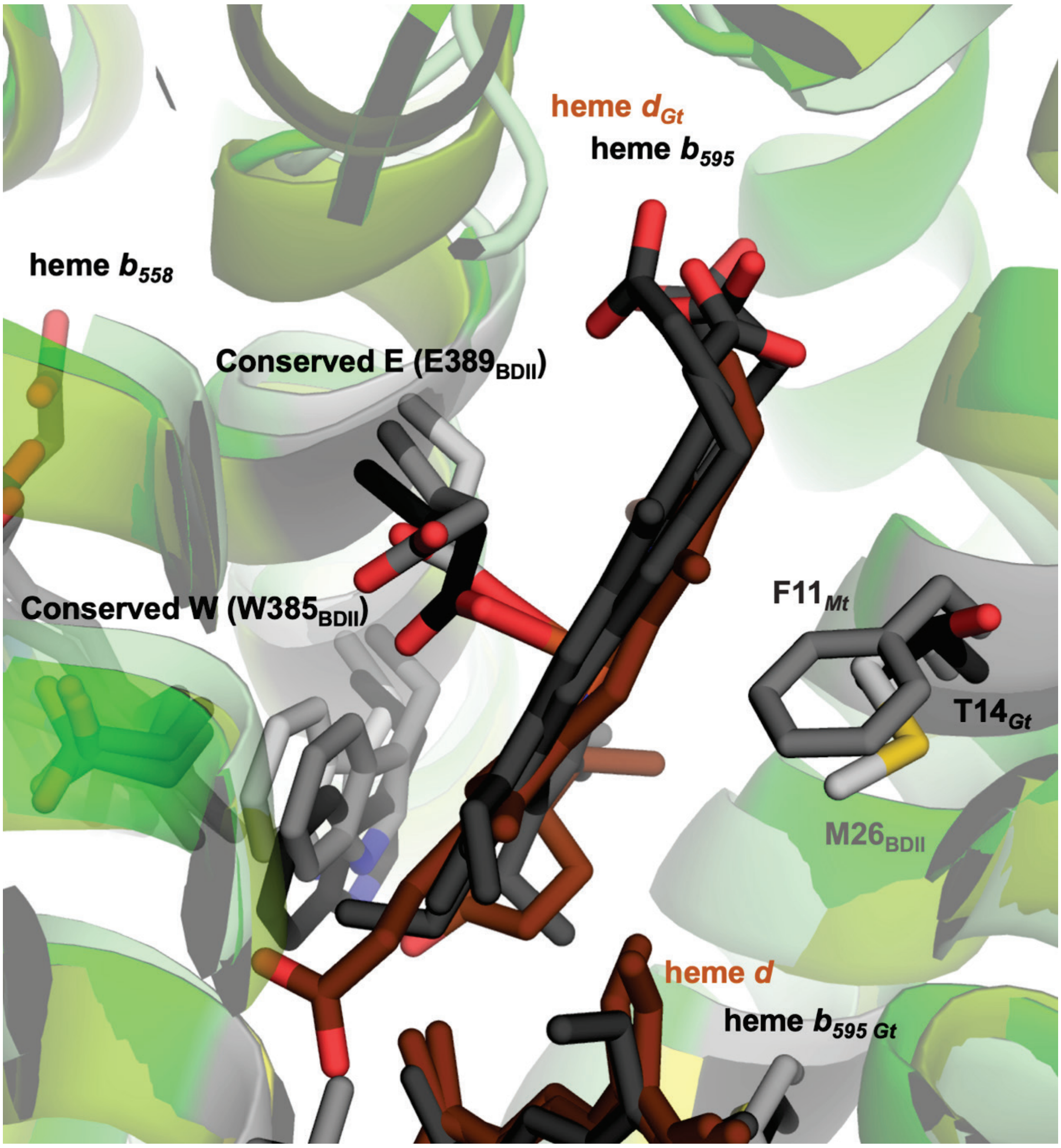


**Figure S7. Ligands and environment of heme $b^{595}$.** Superposition of the *bd*-II oxidase from *M. smegmatis* (light gray), *bd* oxidase from *M. tuberculosis* (dark gray) and the cyt *bd* from *G. thermodenitrificans* (black) demonstrating conserved ligands and different residues in the heme $b^{595}$ environment. Numbering of conserved residues is by *bd*-II from *M. smegmatis.*

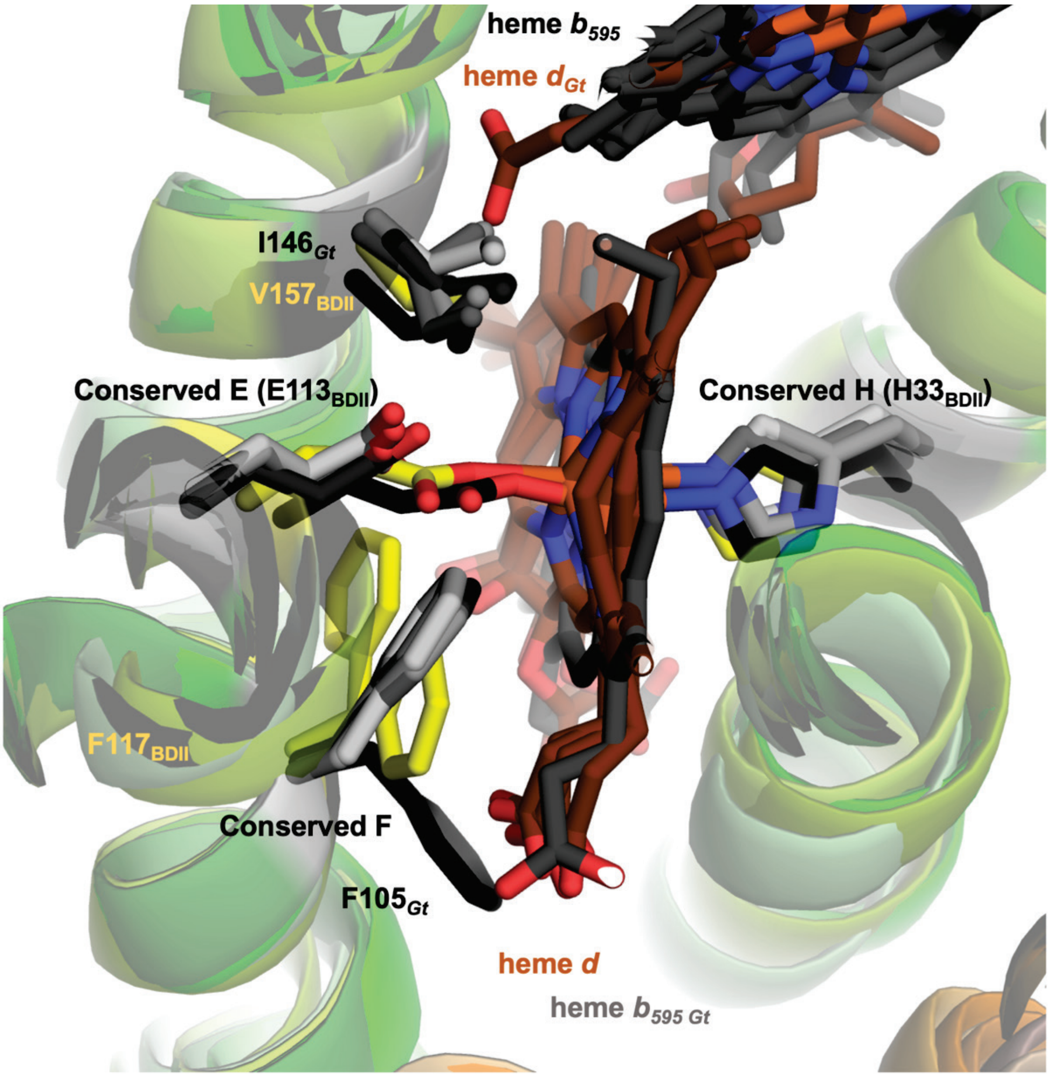


**Figure S8. Ligands and environment of heme *d*.** Superposition of the *bd*-II oxidase from *M. smegmatis* with other known *bd* oxidase structures. Different observed conformation of conserved F residue (yellow: F117 from *bd*-II oxidase from *M. smegmatis* in two conformations) near heme *d* possibly involved in mechanism of activity.

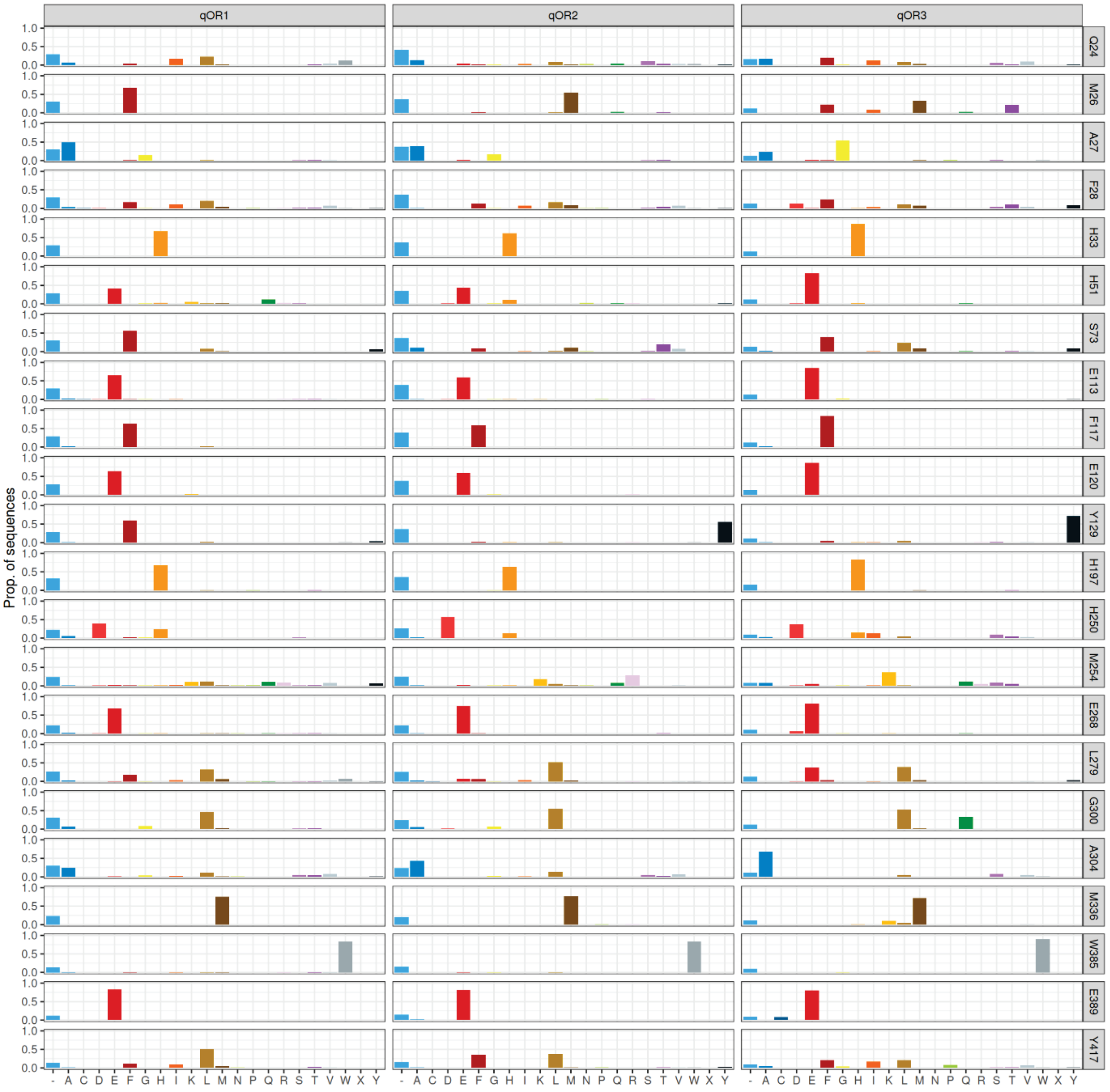


**Figure S9.** Amino acid distribution at selected positions of the *M. smegmatis* AppC and AppB. **A)** AppC. The alignment was made from 3023 sequences resulting from a clustering of all AppC sequences at 60% identity threshold.

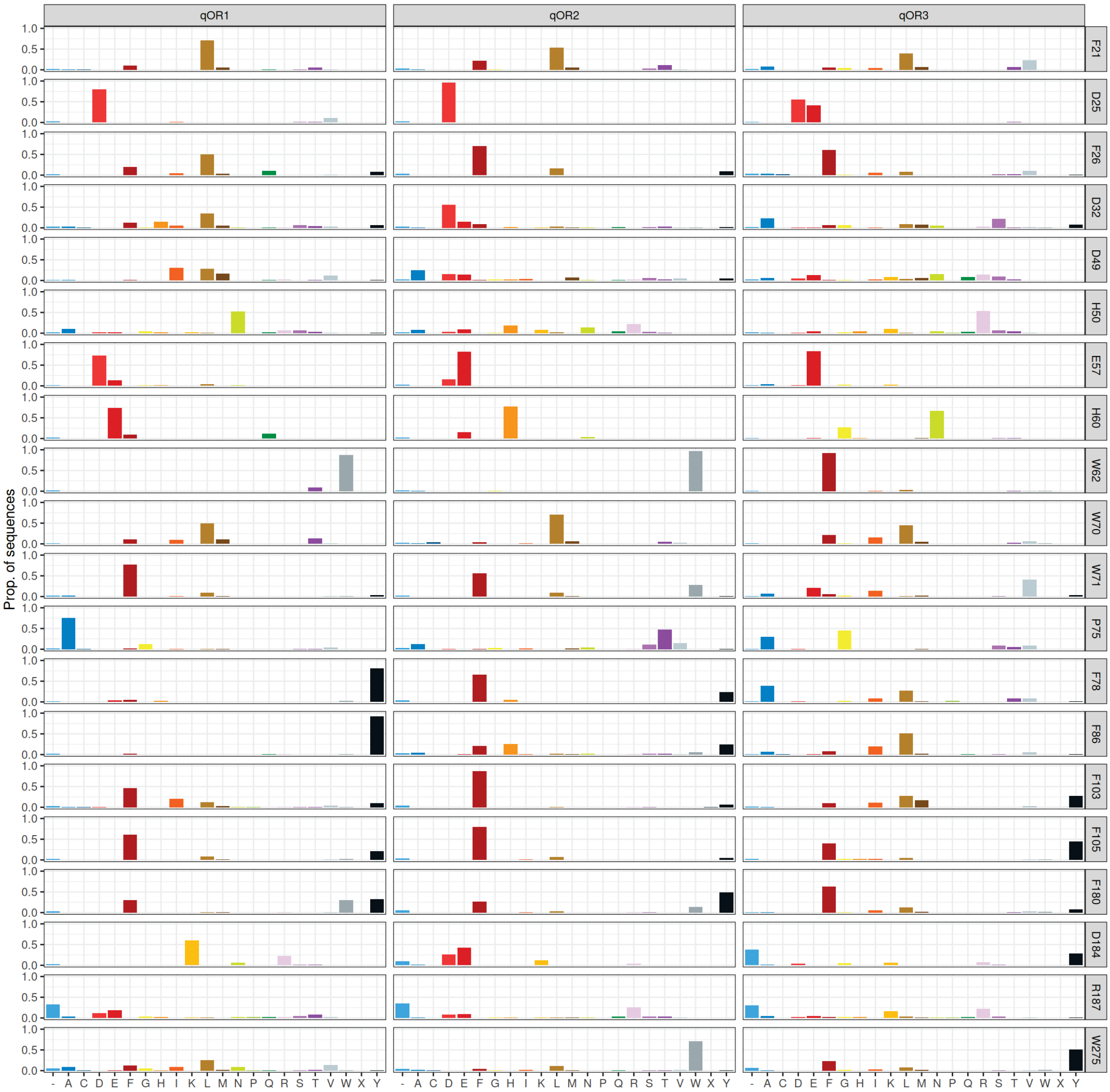


**Figure S9 (contd).** Amino acid distribution at selected positions of the *M. smegmatis* AppC and AppB. B) AppB. The alignment was made from 2007 AppB sequences sitting within five genes from an AppC from which the classification was drawn. The "-" character denotes gaps from proteins that did not align at this position.

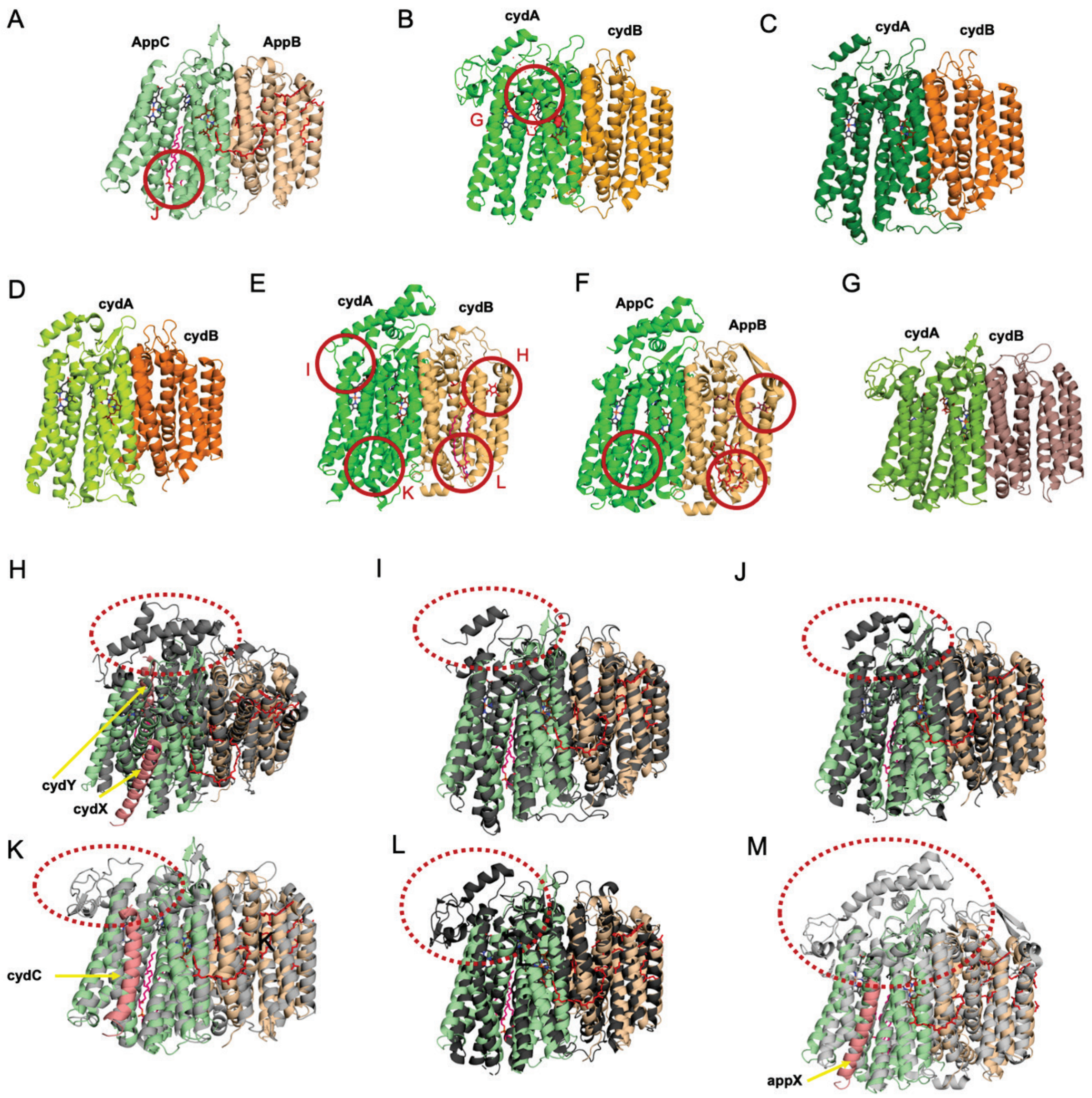


**Figure S10. Comparison of previous cyt *bd* structures. A)** Structure of *bd*-II from *M. smegmatis* (PDB ID: 9R2G). **B)** Structure of the *bd* from *M. tuberculosis* (PDB ID: 7NKZ[7]). **C)** Structure of the cyt *bd*-I from *M. smegmatis* (PDB: 7D5I[8]). **D)** Structure of the cyt *bd*-I oxidase from *C. glutamicum* (PDB ID: 8B4O[9]). **E)** Structure of cyt *bd*-I from *E. coli* (PDB ID: 6RX4[10]). **F)** Structure of cyt *bd*-II from *E. coli* (PDB ID: 7OY2[11]). **G)** Structure of cyt *bd* from *G. thermodenitrificans* (PDB ID: 5DOQ[12]). Individual subunits are differentiated by colour and marked. Red circles mark position of a ligand binding sites with a corresponding letter referring to a window in which the individual binding site is shown in detail. Superposition of the *bd*-II oxidase from *M. smegmatis* (in color) and **H)** cyt *bd*-I from *E. coli.* Extra subunits in pink, **I)** cyt *bd*-I from *M. smegmatis,* **J)** cyt *bd*-I from *C. glutamicum,* **K)** cyt *bd* from *G. thermodenitrificans.* Extra subunit in pink. **L)** cyt *bd* from *M. tuberculosis,* **M)** cyt *bd*-II from *E. coli.* (PDB ID: 7OSE) Red dotted circle marks the Q-loop region.

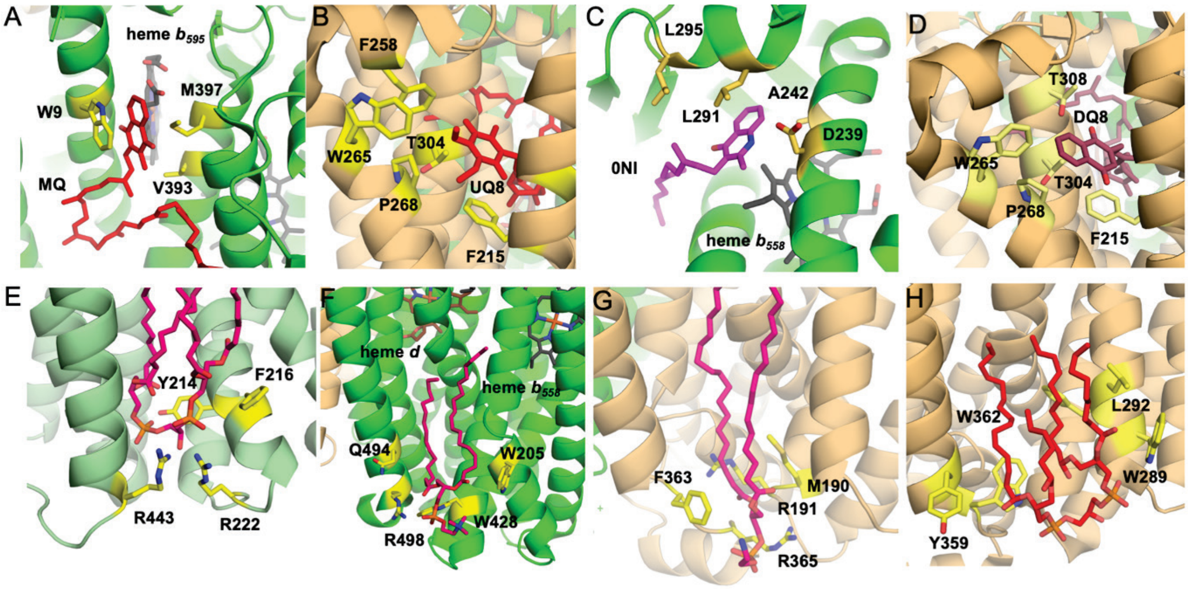


**Figure S11. Ligand binding sites. A)** Extra menaquinone (red) binding site near heme $b^{595}$ in the *M. tuberculosis* cyt *bd* structure. **B)** Extra ubiquinone (red) binding site in subunit cydB of *E. coli bd*-II structure (PDB ID:7OSE). **C)** Aurachin (magenta) bound at the Q-loop position near heme $b_{558}$ of *E. coli bd-II* structure (PDB ID:7OSE). **D)** Extra quinone (red) binding site in subunit appB of *E. coli bd-II* structure (PDB ID:7OY2). **E)** Lipid (pink) binding site in subunit AppC in *bd*-II oxidase from *M. smegmatis.* **F)** Lipid (pink) binding site in subunit AppC in *bd*-II oxidase from *E. coli* (PDB ID:7OY2). **G)** Lipid (pink) binding site in subunit AppB in *bd*-II oxidase from *E. coli* (PDB ID:7OY2). **H)** Lipid (red) binding site in subunit AppB in *bd*-II oxidase from *E. coli* (PDB ID:7OY2).

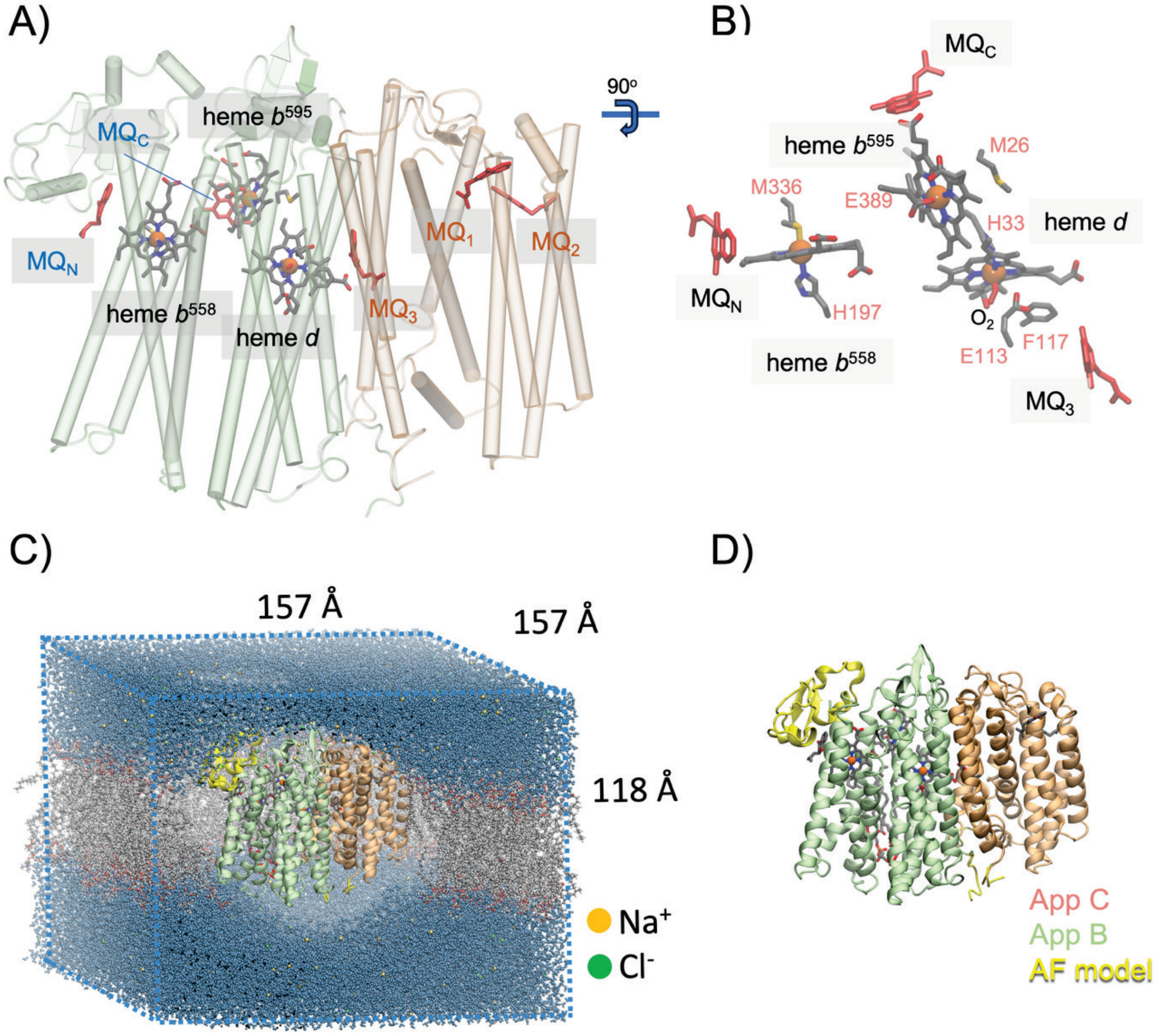


**Figure S12. MD setup of *Ms bd*-II. A)** *bd*-II oxidase model from *M. smegmatis* (PDB ID: 9R2G) with menaquinone (MQ) modelled in Q-loop ($MQ_N$), near heme $b^{558}$ ($MQ_C$) and in AppB ($MQ_1$, $MQ_2$ and $MQ_3$). For clarity only the MQ head is shown, but simulations include the complete MQ-10 isoprenoid tail. **B)** Heme cofactors with modelled ligands. The figure shows $O_2$ modelled in heme-*d* as well as the nearby $E113^C$ and the $F117^C$ observed in two conformations in the cryo-EM map. **C)** The MD simulation system embedded in a water/POPC/POPE/PI/CDL/MQ membrane box comprising around 300,000 atoms. **D)** Model of the unresolved structure based on combination of cryo-EM data and Alphafold[13].

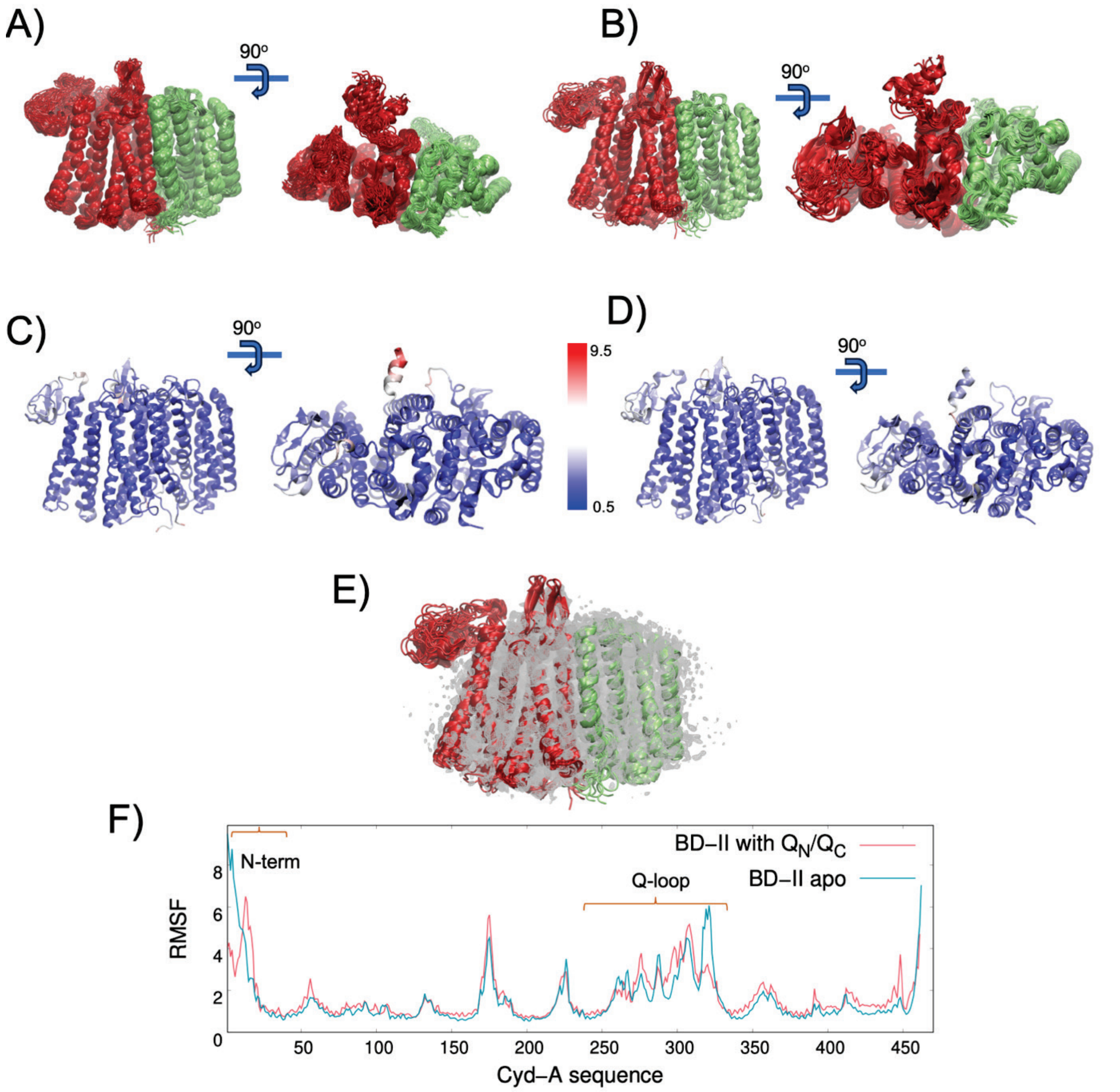


**Figure S13. MD analysis of *Ms bd*-II with and without MQ in the Q-loop.** MD snapshots showing the dynamics of the Q-loop and N-term from AppC for **A, C)** in the *apo* simulation (simulation S5/S6, Table S5) and **B, D**) for the MQ simulated in the Q-loop (simulation S1/S2, Table S5). **E)** Comparison of the cryo-EM map and 50 snapshots extracted from the 1 μs MD simulation of *bd*-II. **C)** RMSF per residue from the 1 μs MD simulation coloured for the apo simulation and **D)** for the MQ in the Q-loop. **F)** Comparison of the RMSF per residue. The large RMSF (at N-term and Q-loop) corresponds to regions not observed in the cryo-EM maps.

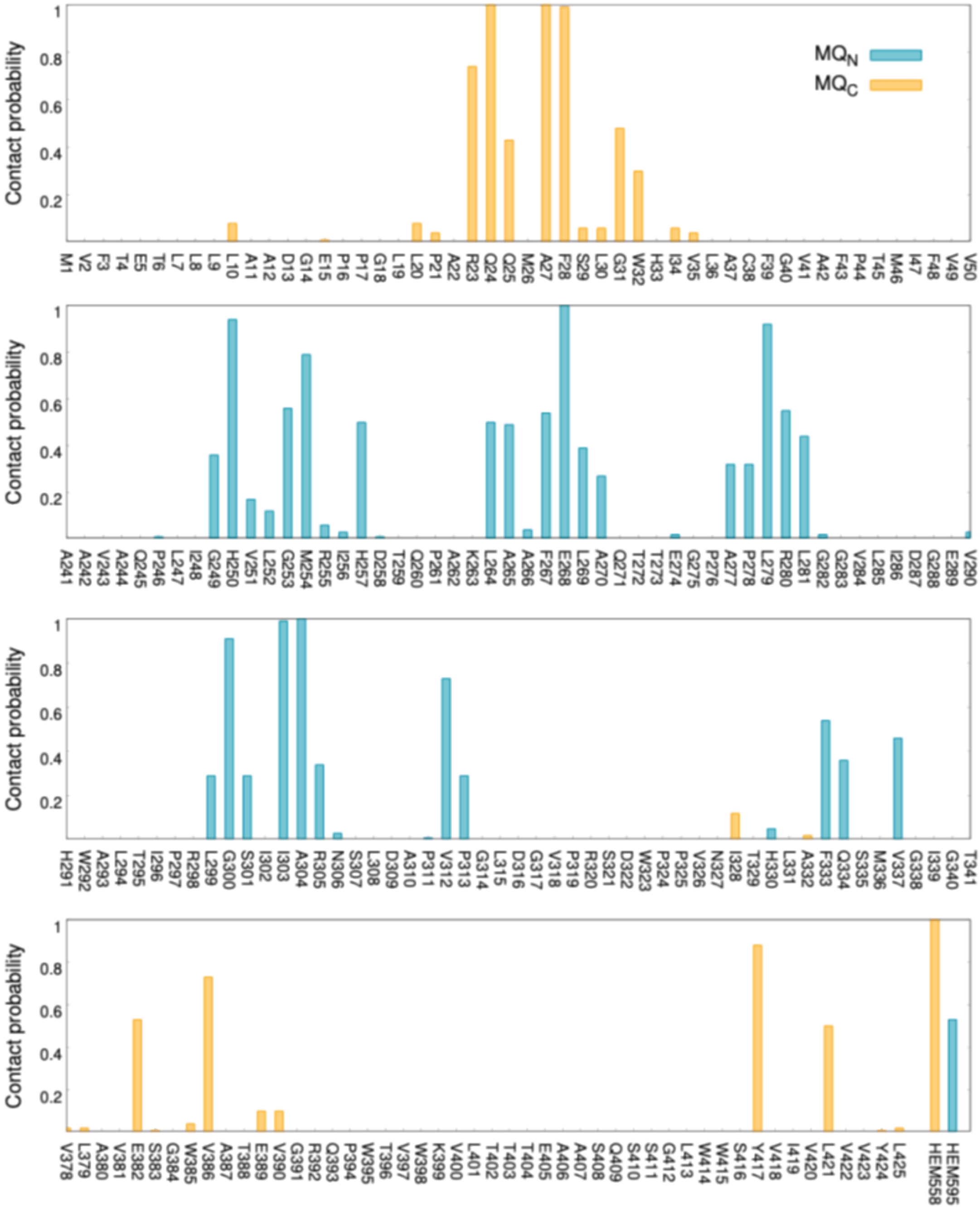


**Figure S14. MQ interactions in the Q-loop.** Statistics of contacts of the MQ headgroup with specific residues in the Q-loop region (AppC). See also Figure 5 of the main text. Residues within 6 Å of MQ were selected for the analysis. Statistical averages and normalization are computed based on 2x2000 frames of 2μs simulation data (simulations S1/S2, Table S5).

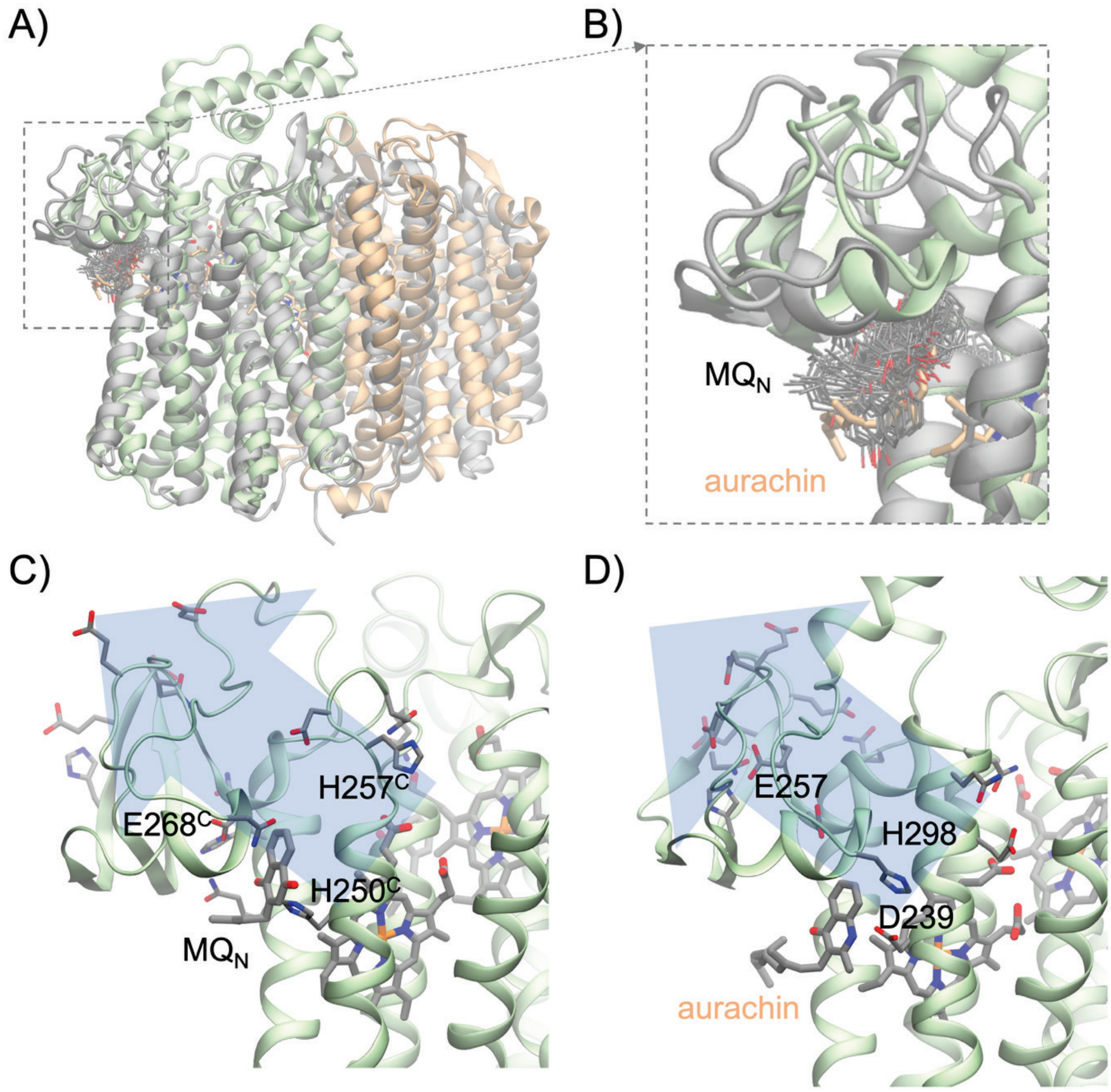


**Figure S15.** Overview of titratable residues in the Q-loop and comparison with aurachin inhibitor binding in *E. coli bd*-II (PDB ID: 7ose[14]). **A)** Overlap of $MQ_N$ dynamics with the position of aurachin in *E. coli bd*-II and **B)** closeup of the Q-loop region. **C, D**) comparison of the conserved residues interacting with $MQ_N$ ($E268^C$ and $H250^C$) and aurachin in *E. coli* (E257 and D239). The Q-loop contains several carboxylic groups that could be support oxidation-coupled proton transfer reactions.

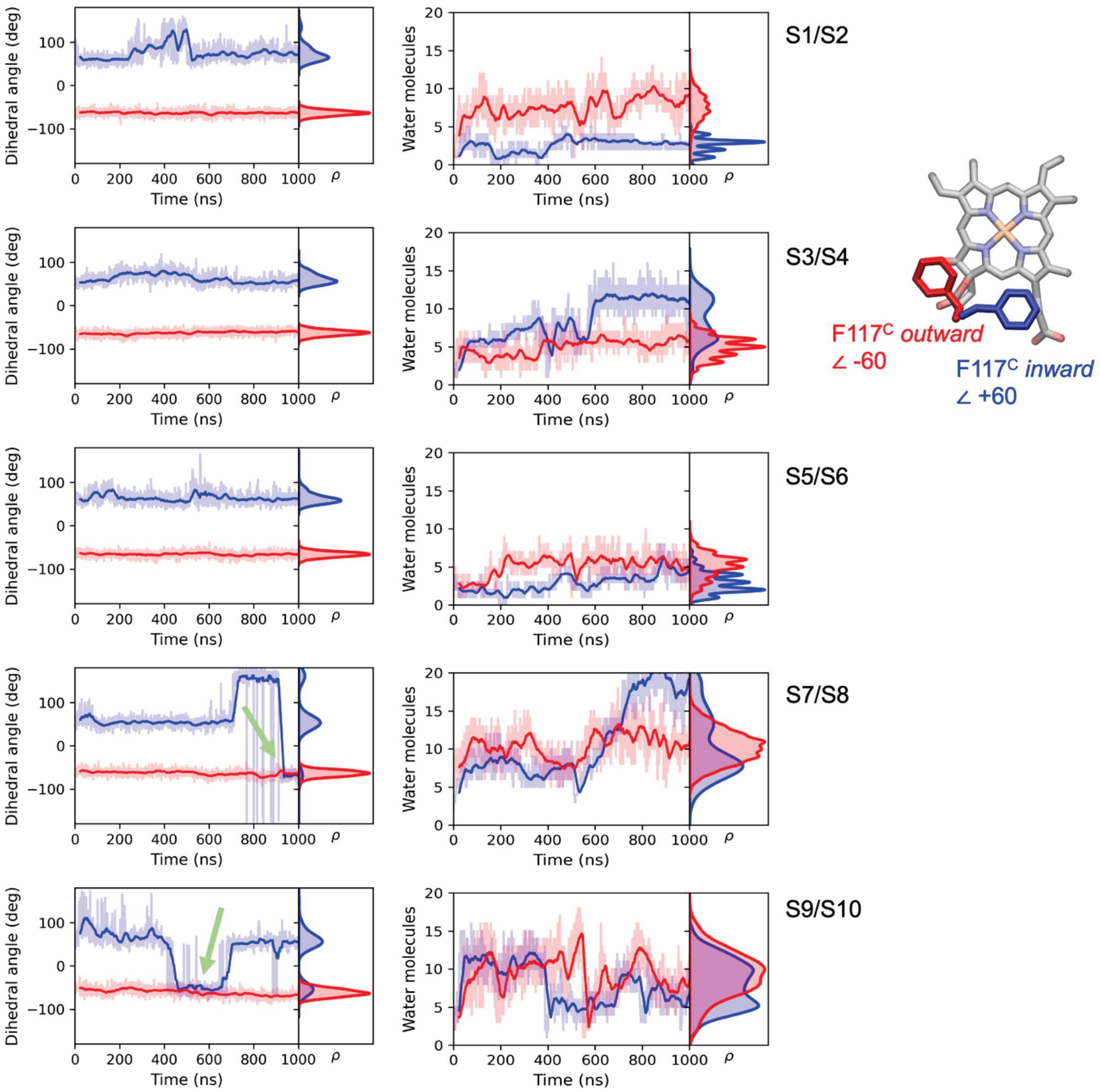


**Figure S16**. Dynamics of F117$^{C}$ (*left*) and hydration level (*right*) from MD simulations in different ligand and protonation states. Left: sampling of "*inward*" (blue) and "*outward*" (red) configurations of F117$^{C}$ (see main text). The F117$^{C}$ configurations remain stable in simulations S1-S6, when E113$^{C}$ is ligated to heme *d*. In the $O_2$ bound heme-*d*, F117$^{C}$ is more dynamic and flips to the "*outward*" conformation (green arrow). This effect is more prominent when E113$^{C}$ is protonated (simulation S9/S10) than for deprotonated state of E113$^{C}$ (simulation S7/S8). *Right*: Hydration near the Fe of heme *d* from MD simulations with different heme *d* ligands and protonation states. The "*outward*" configuration of F117$^{C}$ favours hydration towards the heme *d* when E120$^{C}$ is protonated (simulation S1/S2), and dehydration when E120$^{C}$ is deprotonated (simulations S3/S4).

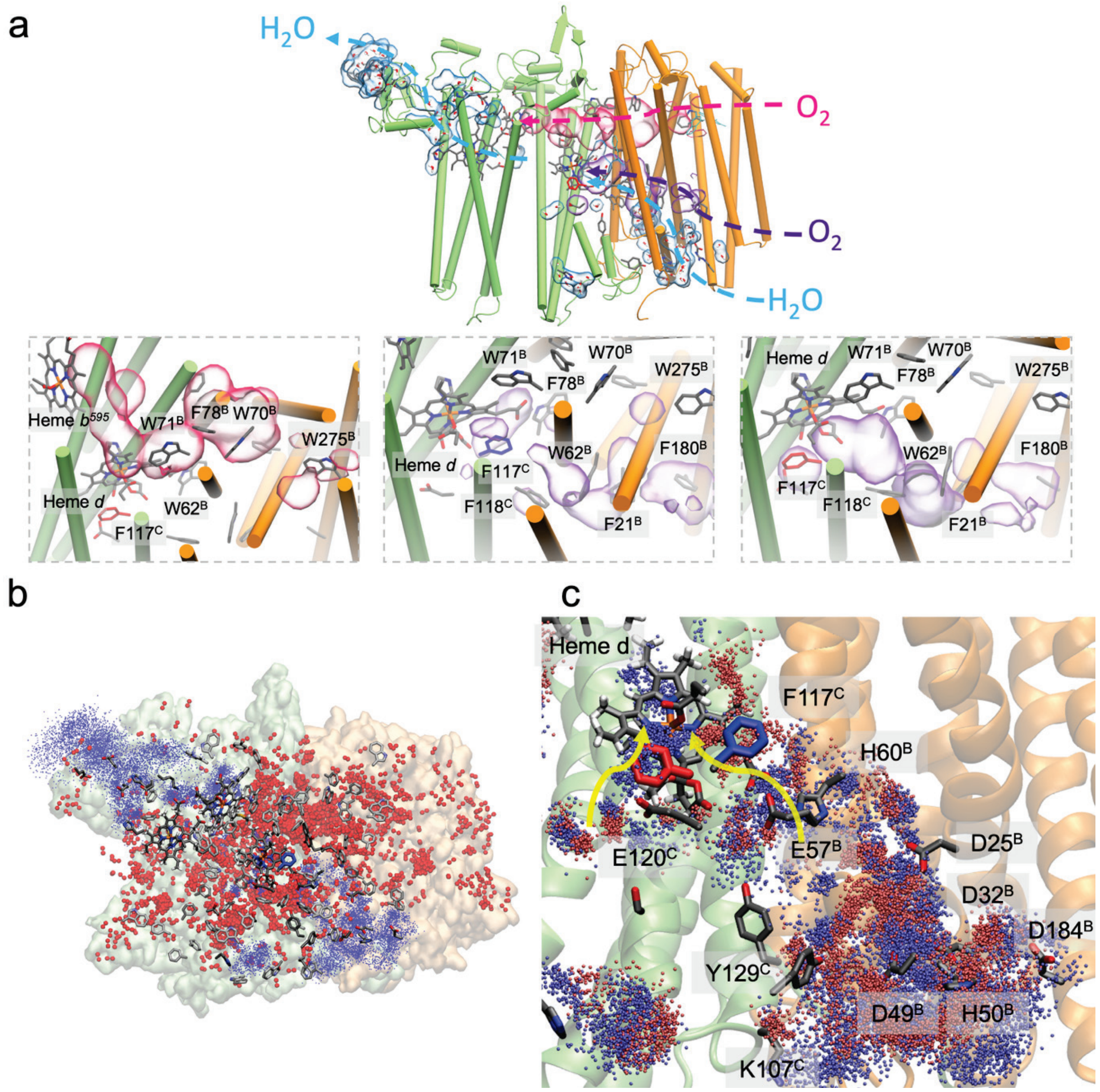


**Figure S17**. Overview of hydration and $O_2$ accessibility from MD simulations for **A)** Top: Two putative dioxygen channels form around heme *d*: in pink the $O_2$ density between heme *d* and heme $b^{595}$ and in purple, $O_2$ pathway via $W62^B$. Density of Proton pathways are depicted in blue and water molecules are represented with sticks for one snapshot. (Bottom figure) Detailed $O_2$ access and relevant residues involved in the paths. Depending on the $F117^C$ orientation, $O_2$ accessibility could be interrupted ($F117^C$ *inward*) or with more access to heme *d*-Fe ($F117^C$ *outward*). **B)** Overlap of water and $O_2$ molecules from 500 ns MD simulations. Water oxygen is represented in small blue spheres while $O_2$ molecules are represented in large red spheres, (see Fig. S16 with water count per simulation). **C)** Comparison of water hydration in both $F117^C$ conformations shows more water accessibility via $E57^B$ when $F117^C$ is facing *outward* (red spheres) and more hydration via $E120^C$ when $F117^C$ is facing *inward* (blue spheres). However, the water channel via $E120^C$ is not connected to the bulk, it is rather connected via $Y129^C$ to AppB water exit/entrance.

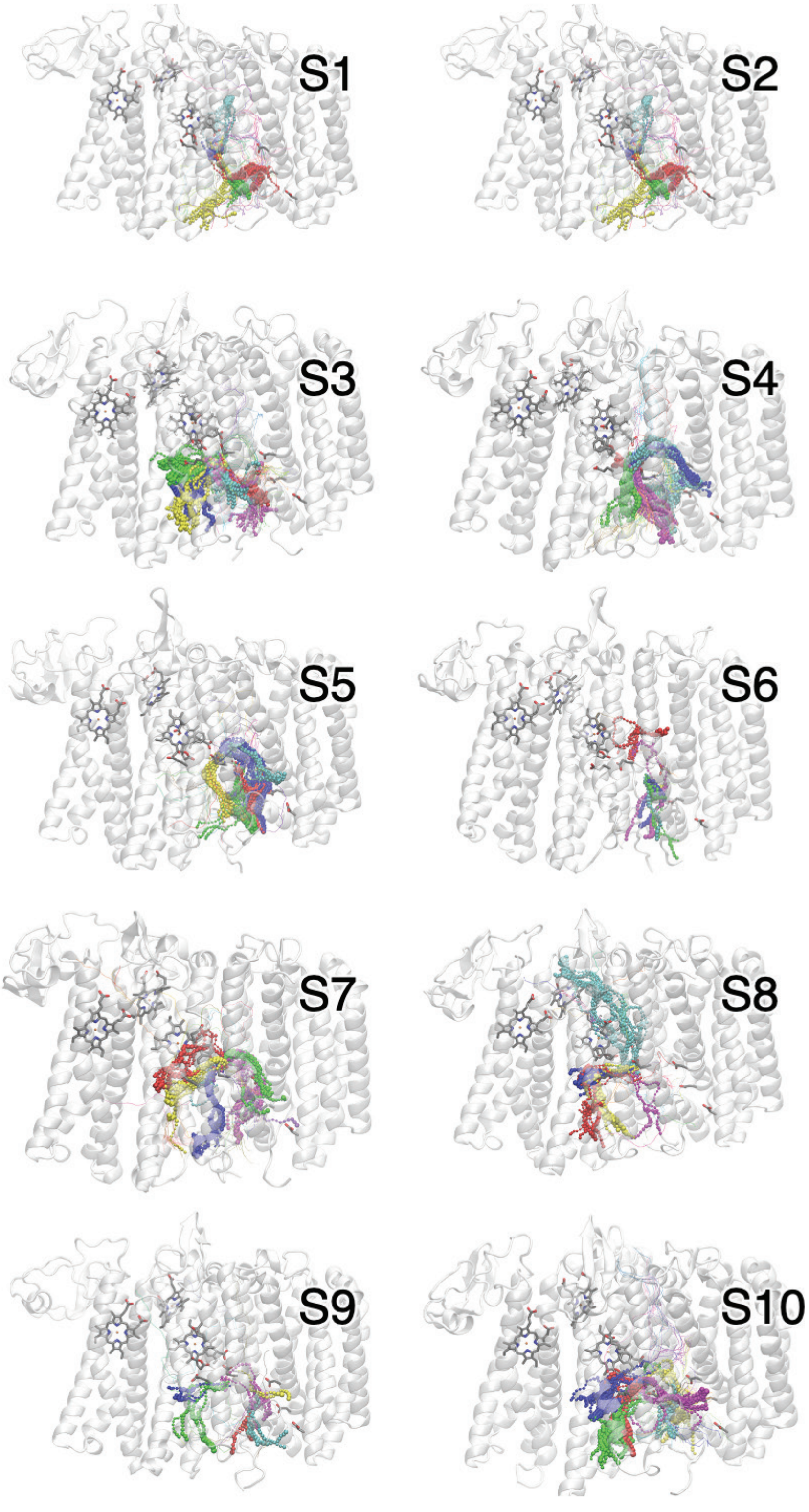


**Figure S18**. Putative proton pathways obtained from MD simulations analyzed using CAVER[15]. The figure shows the five most prominent proton pathways in CPK representation, while the other pathways are shown as lines. The figure highlights also residues E113, E120 and F117 of AppC and E57, D25, D32 and D184 of AppB. The pathways correlate with the position of water molecules observed in MD. The pathways are also sensitive to the orientation of F117 and the protonation states of E113 and E120. $MQ_3$ favors hydration between $E57^B$ and $E113^C$, whereas without $MQ_3$ (simulations S5/S6), the region remains dry.

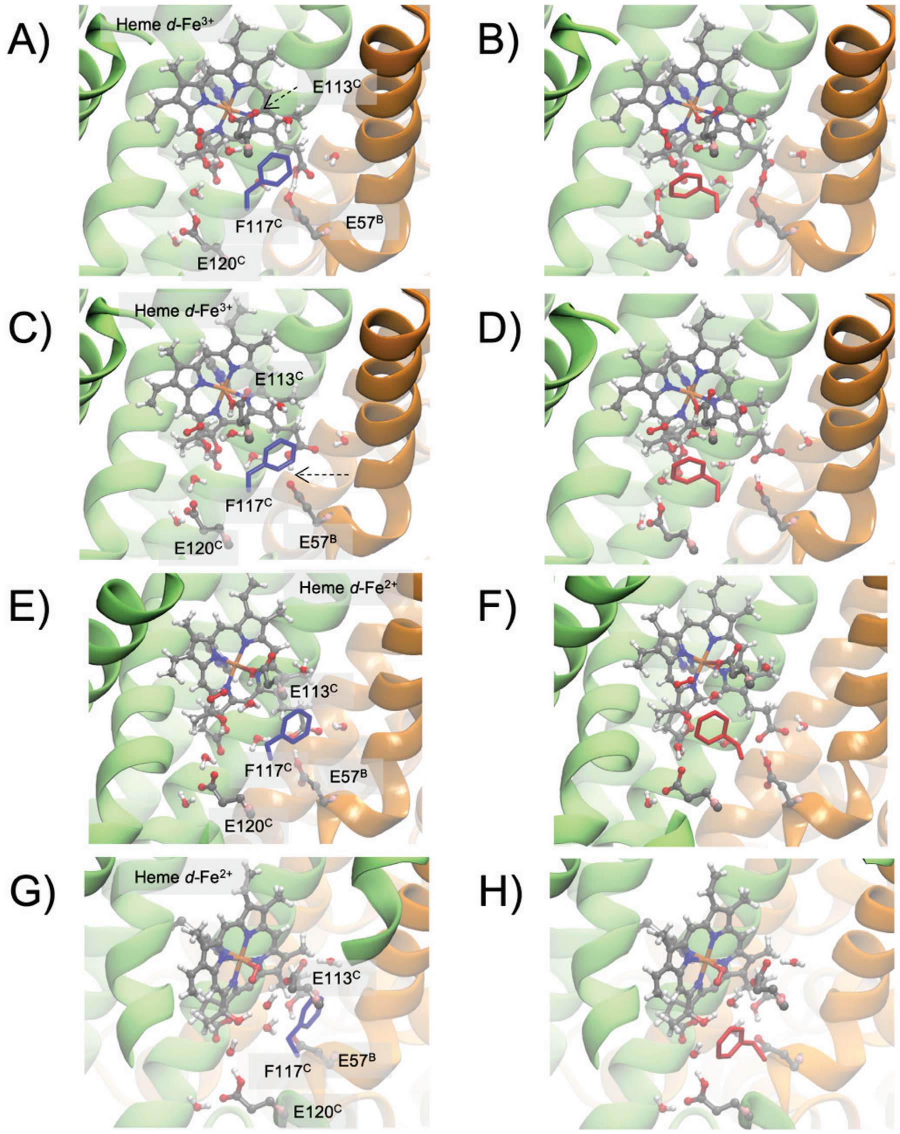


**Figure S19**. Comparison of QM/MM models of heme *d* with different ligands and oxidation state with the cryo-EM data. **A)** Heme *d* ($Fe^{3+}$) with $E113^C$ deprotonated and $F117^C$ in the *inward* and **B)** *outward* conformations. **C)** Heme *d* ($Fe^{3+}$) with $E113^C$ protonated and $F117^C$ in the *inward* and **D)** *outward* conformations. **E)** Heme *d* ($Fe^{2+}$) with $O_2$ bound and $F117^C$ in the *inward* and **F)** *outward* conformations.

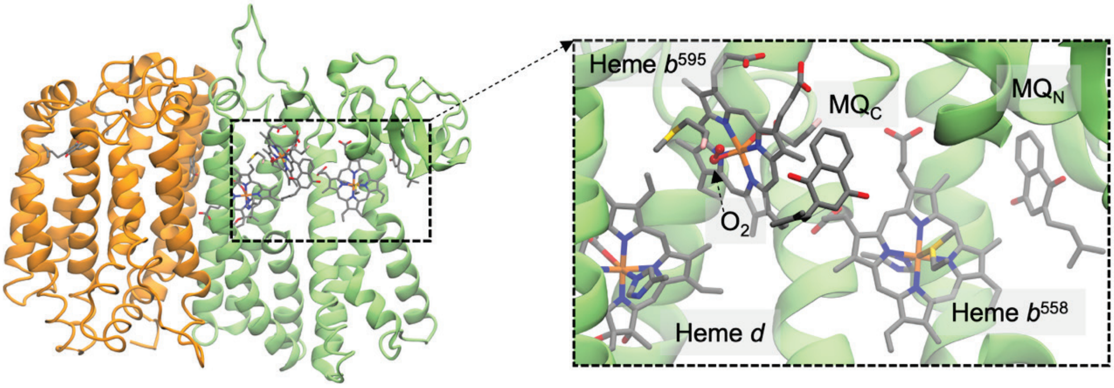


**Figure S20**. QM/MM model of heme $b^{595}$ with bound $O_2$. The distance between Fe-$O_2$ is 1.9 Å in the low spin ($S$=0) $Fe^{2+}$-$O_2$ configuration. The QM/MM model was optimised at the B3LYP-D3/Fe def2-TZVP/def2-TZVP level with the QM region comprised heme $b^{595}$, the $O_2$, E389$^{C}$, M26$^{C}$ (see *Methods*).

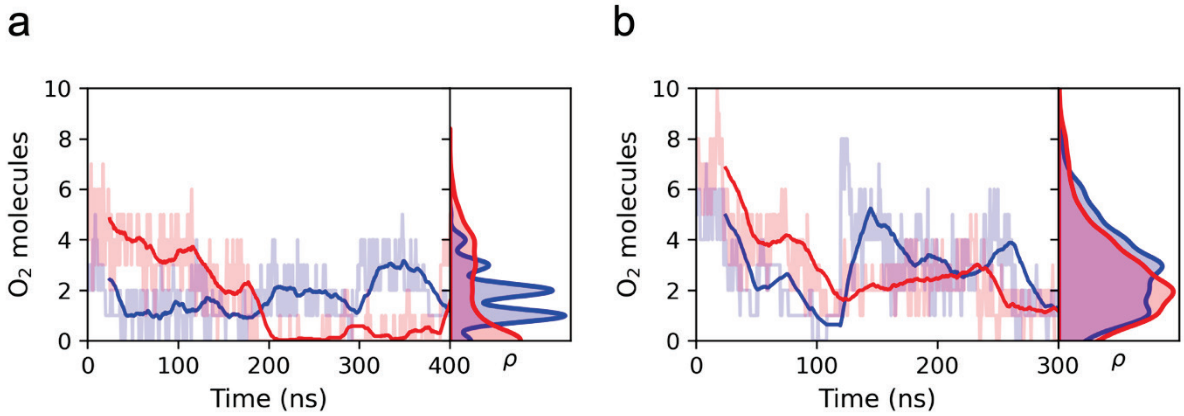


**Figure S21**. Diffusion of $O_2$ near heme $d$ (within 9 Å) with anionic E113$^{C}$ (*left*) and neutral E113$^{C}$ (*right*) (from simulations S7/S8 and S9/10, respectively). The figure shows number of $O_2$ molecules during the last 300 ns of the MD simulations in the "*inward*" (blue) and "*outward*" (red) configurations of F117$^{C}$, with the simulations initiated from the same initial positions of $O_2$ (see Extended Methods). More oxygen molecules accumulate near heme $d$, when E113$^{C}$ is neutral and F117$^{C}$ is in the "*inward*" configuration.

**Table S1.** Actinomycetota species with only qOR-2 *bd*.

| class | order | family | % species with only qOR2 |
|---|---|---|---|
| Actinomycetes | Propionibacteriales | Nocardioidaceae | 11 |
| Thermoleophilia | Solirubrobacterales | sep-70 | 53 |
| Actinomycetes | Mycobacteriales | Frankiaceae | 67 |
| Actinomycetes | Streptosporangiales | Streptosporangiaceae | 6 |
| Actinomycetes | Mycobacteriales | Pseudonocardiaceae | 5 |
| Acidimicrobiia | Acidimicrobiales | SZUA-35 | 40 |
| Acidimicrobiia | Acidimicrobiales | Ilumatobacteraceae | 7 |
| Acidimicrobiia | Acidimicrobiales | SHLQ01 | 70 |
| Thermoleophilia | Gaiellales | Gaiellaceae | 11 |
| Thermoleophilia | Solirubrobacterales | Thermoleophilaceae | 35 |
| Acidimicrobiia | IMCC26256 | PALSA-555 | 91 |
| Actinomycetes | Mycobacteriales | Mycobacteriaceae | 1 |
| Actinomycetes | Mycobacteriales | Geodermatophilaceae | 8 |
| Thermoleophilia | Gaiellales | F1-60-MAGs149 | 50 |
| Thermoleophilia | Miltoncostaeales | Miltoncostaeaceae | 19 |
| Actinomycetes | Mycobacteriales | Micromonosporaceae | 2 |

**Table S2.** Data collection and processing of cryo-EM data.

| Data collection, processing | *Ms bd*-II |
|---|---|
| PDB code | 9R2G |
| Voltage (kV) | 300 |
| Magnification | 130 000 |
| Electron exposure ($e^{-}$/Å$^{2}$) | 93.5 |
| Pixel size (Å) | 0.650 |
| Defocus range (µm) | -2.2 to -0.6 |
| Defocus step (µm) | -0.2 |
| Symmetry imposed | None C1 |
| Initial particle images (number) | 7 907 011 |
| Final particle images (number) | 200 084 |
| FSC threshold | 0.143 |
| Map resolution (Å) | 2.8 |
| **Refinement** | |
| CC (mask) | 0.87 |
| Resolution estimates (Å) | |
| d 99 | |
| Masked | 3.6 |
| Unmasked | 3.6 |
| d FSC model, 0/0.143/0.5 (Å) | |
| Masked | 2.8/2.8/3.0 |
| Unmasked | 2.8/2.9/3.1 |
| Model composition | |
| Protein residues | 707 |
| Non-hydrogen atoms | 5774 |
| Ligands | 7 |
| Water | 7 |
| B factors (Å$^{2}$) | |
| Min/max/mean | |
| Protein residues | 64.43/191.82/101.94 |
| Ligands | 69.11/161.38/98.06 |
| water | 86.55/113.33/99.03 |
| **Validation** MolProbity | |
| Ramachandran plot | |
| Favoured (%) | 97.72 |
| Allowed (%) | 2.14 |
| Disallowed (%) | 0.14 |

**Table S3.** List of primers used for quantitative real-time PCR.

| Gene | | Sequence (5′→3′) | $T_m$(°C) |
|---|---|---|---|
| *cydA* | Forward | TTC CTC ACC GCA GGT GTT TT | 57.3 |
| | Reverse | TGC TGC TCG AAC ATC AGC TT | 57.3 |
| *cydB* | Forward | GTT CCC GGA TCT CAT CCC CT | 61.3 |
| | Reverse | CGC TTG CTG AAC ACC CAG TA | 59.4 |
| *appC* | Forward | CAT CTG GCG TTC CAG TCG AT | 59.4 |
| | Reverse | TGA CGT CCT ACT TCG GTT GC | 59.4 |
| *appB* | Forward | AGT TTC GCG TTC CGC AAG TA | 57.3 |
| | Reverse | CGA CAG TCC CCA GGA AGA AC | 61.3 |
| *sigA* | Forward | CTC AAC GCC GAA GAA GAG GT | 59.4 |
| | Reverse | CTG GAT GAG GTC GAG GAA CG | 61.4 |

**Table S4**. List of relevant amino acid and their corresponding numbers in selected cyt *bd*s species.

| ***Ec bd*-I** | ***Ms bd-II*** | ***Ms bd*-I** | ***Gt bd*** |
|---|---|---|---|
| Phe-104[A] | Phe-117[C] | Phe-103[A] | Phe-105[A] |
| Glu-99[A] | Glu-113[C] | Glu-98[A] | Glu-101[A] |
| Glu-107[A] | Glu-120[C] | Glu-106[A] | Glu-108[A] |
| His-186[A] | His-197[C] | His-185[A] | His-186[A] |
| Met-393[A] | Met-336[C] | Met-346[A] | Met-325[A] |
| Trp-441[A] | Trp-385[C] | Trp-394[A] | Trp-374[A] |
| Glu-445[A] | Glu-389[C] | Glu-398[A] | Glu-378[A] |
| His-19[A] | His-33[C] | His-18[A] | His-21[A] |
| Phe-12[A] | Met-26[C] | Phe-11[A] | Thr-14[A] |
| Asp-58[B] | Glu-57[B] | Asp-62[B] | Glu-57[B] |
| Trp-63[B] | Trp-62[B] | Trp-67[B] | Phe-62[B] |

**Table S5. List of molecular dynamics (MD) simulations**. The MD system with around 280,000 atoms (see *Methods*). Heme $b^{558}$, heme $b^{595}$ and heme d *d* ($Fe^{II}$-$O_2$), were modelled in their oxidized states, whereas the MQ was modelled as menaquinol in $MQ_N$ and $MQ_C$, and as menaquinone in $MQ_1$, $MQ_2$, and $MQ_3$. [a,b] Resides modeled in their neutral/anionic states, respecitvely.

| **Simulation** | **$MQ_N$** | **$MQ_C$** | **heme *d* ligand** | **F117** | **Length (ns)** | **Protonation[a,b]** |
|---|---|---|---|---|---|---|
| **S1** | $QH_2$ | $QH_2$ | E113 | *inward* | 2 x 500 | E120[a], E57[b] |
| **S2** | $QH_2$ | $QH_2$ | E113 | *outward* | 2 x 500 | E120[a], E57[b] |
| **S3** | $QH_2$ | $QH_2$ | E113 | *inward* | 2 x 500 | E57[b] |
| **S4** | $QH_2$ | $Q_{o1a}$ | E113 | *outward* | 2 x 500 | E57[b] |
| **S5** | - | - | E113 | *inward* | 2 x 500 | E120[a], E57[b] |
| **S6** | - | - | E113 | *outward* | 2 x 500 | E120[a], E57[b] |
| **S7** | $QH_2$ | $QH_2$ | $O_2$ | inward | 2 x 500 | E120[a], E57[b] |
| **S8** | $QH_2$ | $QH_2$ | $O_2$ | *outward* | 2 x 500 | E120[a], E57[b] |
| **S9** | $QH_2$ | $QH_2$ | $O_2$ | *inward* | 2 x 500 | E113[a], E120[a], E57[b] |
| **S10** | $QH_2$ | $QH_2$ | $O_2$ | *outward* | 2 x 500 | E113[a], E120[a], E57[b] |
| | | | | **Total:** | **10.0 μs** | |

**Table S6**. Heme *d* ligands and bond distances from QM/MM models. $E113^{-}$ and $E113^{0}$ refer to anionic and neutral forms of E113, respectively.

| Model | Fe-His(NE2) (Å) | Fe-Glu(OE2) (Å) | $Fe-O_2$ (Å) | $F117^{C}$ |
|---|---|---|---|---|
| **heme *d* ($Fe^{3+}$) – $E113^{-}$** | 2.75 | 2.10 | - | *inward* |
| **heme *d* ($Fe^{3+}$) – $E113^{-}$** | 2.53 | 2.25 | - | *outward* |
| **heme *d* ($Fe^{3+}$) – $E113^{0}$** | 2.22 | 3.06 | - | *inward* |
| **heme *d* ($Fe^{3+}$) – $E113^{0}$** | 2.17 | 2.84 | - | *outward* |
| **heme *d* ($Fe^{2+}$) – $E113^{0}$** | 2.22 | 3.05 | - | *inward* |
| **heme *d* ($Fe^{2+}$) – $E113^{0}$** | 2.23 | 2.86 | - | *outward* |
| **heme *d* ($Fe^{2+}$) – $O_2$** | 2.36 | | 2.08 | *inward* |
| **heme *d* ($Fe^{2+}$) – $O_2$** | 2.34 | | 2.10 | *outward* |
